# GH18 family glycoside hydrolase Chitinase A of *Salmonella* facilitates bacterial invasion and survival by modulating host immune responses

**DOI:** 10.1101/2021.07.13.452150

**Authors:** Kasturi Chandra, Dipshikha Chakravortty

## Abstract

*Salmonella* is a facultative intracellular pathogen that has co-evolved with its host and has also developed various strategies to evade the host immune responses. *Salmonella* recruits an array of virulence factors to escape from host defense mechanisms. Previously *chitinase A* (*chiA*) was found to be upregulated in intracellular *Salmonella*. Although studies show that chitinases and chitin binding proteins (CBP) of many human pathogens have a profound role in various aspects of pathogenesis, like adhesion, virulence and immune evasion, the role of chitinase in strict intravacuolar pathogen *Salmonella* has not yet been elucidated. In this study, we deciphered the role of chitinase of *Salmonella* in the pathogenesis of the serovars, Typhimurium and Typhi. Our data propose that ChiA mediated modification of the glycosylation on the epithelial cell surface facilitates the invasion of the pathogen into the epithelial cells. Further we found that ChiA aids in reactive nitrogen species (RNS) and reactive oxygen species (ROS) production in phagocytes, leading to MHCII downregulation followed by suppression of antigen presentation and antibacterial responses. In continuation of the study in animal model *C. elegans*, *Salmonella* Typhi ChiA was found to facilitate attachment to the intestinal epithelium, gut colonization and persistence by downregulating antimicrobial peptides.

## Introduction

*Salmonella* is one of the major foodborne pathogens that causes enteric disturbances in humans and other mammals. Although *Salmonella*-mediated enteric illnesses can be treated, the high occurrences of drug-resistant strains challenge the pathogen eradication. Human gastrointestinal tract is covered with two distinct types of glycan layers-mucin and complex oligosaccharides (glycocalyx) that protects the enterocytes from the environment [1]. To gain access to the enterocytes, an enteric pathogen like *Salmonella* should be able to cleave the mucinous layer. In various human pathogens, glycoside hydrolases such as sialidases, muraminidases, glucosaminidases, pullulanases, GalNAcases etc. are known to facilitate the bacterial attachment to the host cells [2]. GH18 family protein chitinases and chitin binding proteins were also found to be involved in pathogenesis of several human enteric (*Vibrio cholerae, Listeria monocytogenes, Serratia marcescens*) [3–7] and non-enteric pathogens (*Pseudomonas aeruginosa, Legionella pneumophila*) [8–10]. In all these pathogens, commonality of the presence of mucin-rich environment hinted towards a potentially significant role of chitinases and chitin binding proteins in breaching mucosal barrier. *Salmonella* causes infection in the gut mucosal region which also has a protective mucinous layer. A BLAST search revealed that *Salmonella* Typhimurium exochitinase ChiA (encoded by STM14_0022) showed 20-40% identity with the abovementioned pathogenic proteins. Further *Salmonella* Typhi chitinase (ChiA; STY0018) is 98% similar to the *S.* Typhimurium SL1344 *chiA* (STM0022) that was reported to be upregulated ∼12-20 fold in the infected macrophages and ∼4-5 fold in the epithelial cells [11, 12].

We infected epithelial cells and phagocytes with the mutant strain and interestingly we found that the mutant was invasion defective in epithelial cells. *Salmonella* is known to remodel the host cell surface glycans to facilitate invasion in the epithelial cells [13–15], we checked the host cell surface glycan modification by lectin-binding assay. Our data suggest that chitinase aids in glycan remodeling by cleaving the terminal sialic acid (Neu5Ac), and Gal-β1,4-GalNAc, thus making the mannose residues accessible to the bacteria for binding. Further we found that the phagocytes infected with the mutant bacteria produced less antibacterial molecules. Interestingly, the mutants were significantly less virulent, less persistent, and were unable to dampen host antibacterial and immune responses in the *in vivo* infection models. Moreover, in this study we demonstrated a novel role of ChiA in facilitating extra-intestinal colonization of *Salmonella* Typhi in *C. elegans.* Together our data suggest that chitinase A plays a multifaceted role in *Salmonella* pathogenesis ranging from aiding bacterial invasion in epithelial cells, enhancing antibacterial NO production *ex vivo*, to increasing bacterial persistence in the nematodes and regulating cellular and humoral immune responses *in vivo*.

## Results

### Chitinase deletion impairs bacterial invasion in human epithelial cells

Since previous reports suggest that chitinases and chitin-binding proteins (CBPs) were involved in adhesion, invasion and *in vivo* pathogenesis of several human pathogens [3–10] and STM ChiA and STY ChiA are 19-24% identical to these pathogenic proteins (**Fig. S1A**), we made isogenic mutants of *chiA* using lambda red recombinase method [16]. The mutants as well as the trans-complemented strain (STY Δ*chiA:chiA*) did not show any growth difference *in vitro* (**Fig. S1B, S1C**), suggesting that Chitinase A is non-essential for extracellular life of *Salmonella sp*. Upon entering the host and surviving through the acidic stomach environment, *Salmonella* reaches gut epithelium, where SPI1-T3SS effectors induce membrane ruffling in the enterocytes. This facilitates bacterial entry in the epithelial cells and marks the beginning of *Salmonella* infection [17]. Since earlier report suggested that *chiA* of *S*. Typhimurium SL1344 strain (STM0022) was highly upregulated in intracellular bacteria from epithelial cells, we checked bacterial invasion and intracellular proliferation in Caco2 cells. We found that the *chiA* deletion rendered the bacteria less invasive and hyperproliferative in epithelial cells (**Fig. 1A-D, S1D, S1E**). We next checked the expression of SPI1 and SPI2 effector genes in intracellular bacteria as SPI1 effectors facilitate bacterial invasion in epithelial cells and SPI2 effectors are required for intracellular survival and proliferation. Surprisingly we found that SPI1 effectors *invF* and *hilA* were significantly upregulated during the early phase of infection in the Δ*chiA* mutant bacteria, whereas no significant difference was observed in the expression of SPI2 effector *ssaV* after 16 hours of infection (**Fig. 1E-F**), suggesting that the reduced bacterial invasion in the epithelial cells by Δ*chiA* mutant is independent of SPI1 gene expression.

**Fig 1.**
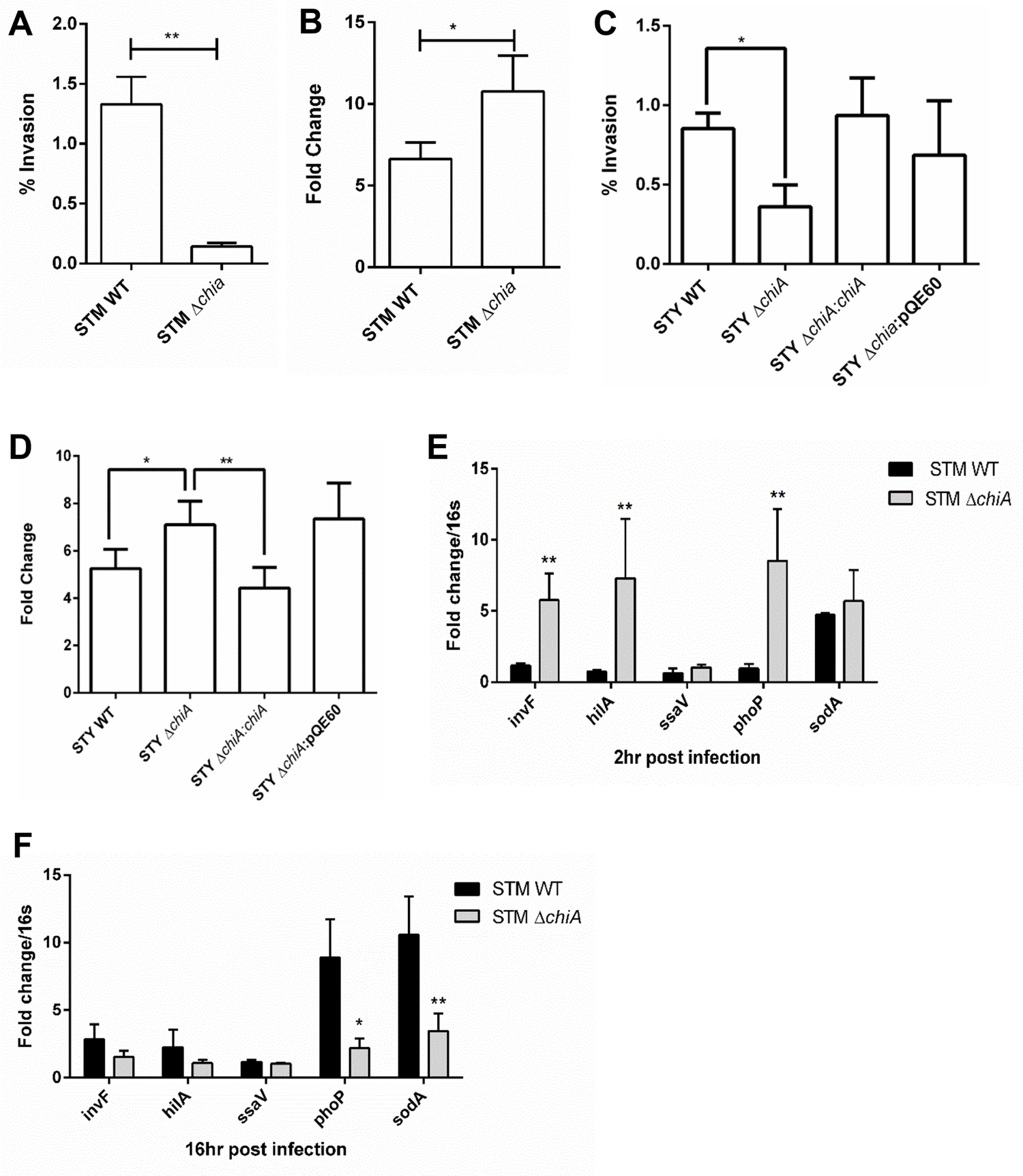
Chitinase deletion impairs bacterial invasion in human epithelial cells. **(A)** % invasion and **(B)** Intracellular proliferation of STM WT and STM Δ*chiA* strains in Caco2 cells by gentamicin protection assay. Data are represented as mean ± SEM of 3 independent experiments (N=3, n=3). Unpaired Student’s t test was used to analyze the data. **(C)** % invasion and **(D)** Intracellular proliferation of STY WT, STY Δ*chiA,* STY Δ*chiA:chiA* and STY Δ*chiA:*pQE60 strains in Caco2 cells by gentamicin protection assay. Data are represented as mean ± SEM of 3 independent experiments (N=3, n=3). One-way ANOVA was used to analyze the data. **(D-E)** RNA expression level of SPI1 and SPI2 genes in from STM WT and STM Δ*chiA* during **(D)** early phase and **(E)** late phase of infection in Caco2 cells. Data are represented as mean ± SEM of 3 independent experiments (N=3, n=3). Two-way ANOVA was used to analyze the data.

### Chitinase A facilitates bacterial entry in epithelial cells by cell surface glycan modification

The intestinal epithelium is covered with mucus and the cells are layered with oligosachharide molecules that forms the glycocalyx. α2-6, α2-3, α1-3, β1-3 or β1-6 linked Glycosylation on the host epithelial cells follow a particular array, such as the outermost glycosylation is N-acetylneuraminic acid (Neu5Ac or sialic acid), followed by galactose, N-acetylgalactosamine (GalNAc), N-acetylglucosamine (GlcNAc) and the innermost moieties mannose and fucose (**Fig. 2A**) [14]. To initiate infection, an enteric pathogen must breach the protective glycocalyx layer. Particularly in the pathogenesis of *S.* Typhi, host cell surface glycoproteins are known to play an important role in typhoid toxin mediated inflammation [18]. Therefore, we checked the abundance of various glycosyl molecules present on Caco2 cells after *Salmonella* infection. Interestingly we observed that after 120 min of infection with the WT bacteria, the host cell surface showed a significant decrease in the abundance of sialylation on the surface glycome, which was not observed for the cells infected with Δ*chiA* mutants, suggesting chitinase is involved in removal of the terminal sialic acids (**Fig. 2B; top panel**). This was further validated by the shift of the Neu5Ac-bound SNA-FITC lectins towards lower abundance in flow cytometry analysis (**Fig. 2C; first column**). Consequently, we observed a significant increase in the abundance of Gal-β1,3-GalNAc on the cells infected with WT bacteria as compared to the cells infected with Δ*chiA* strains (**Fig. 2B; middle panel**), which was further corroborated by a significant shift of Gal-bound PNA-FITC lectins towards higher abundance in flow cytometry analysis (**Fig. 2C; second column**). Finally, we observed a substantial increase in the abundance of mannose-bound concanavalin A-FITC fluorescence on the cell surface of WT bacteria infected cells as compared to the Δ*chiA* mutant infected cells (**Fig. 2B; bottom panel**) that was supported by a significant shift of mannose-bound ConA-FITC lectins towards higher abundance in flow cytometry analysis (**Fig. 2C; third column**). The cell surface glycan bound lectin fluorescence was further quantified which further validated these observations (**Fig. S1F**) Together these data suggested that *Salmonella* chitinase helps in the host cell surface remodeling.

**Fig 2.**
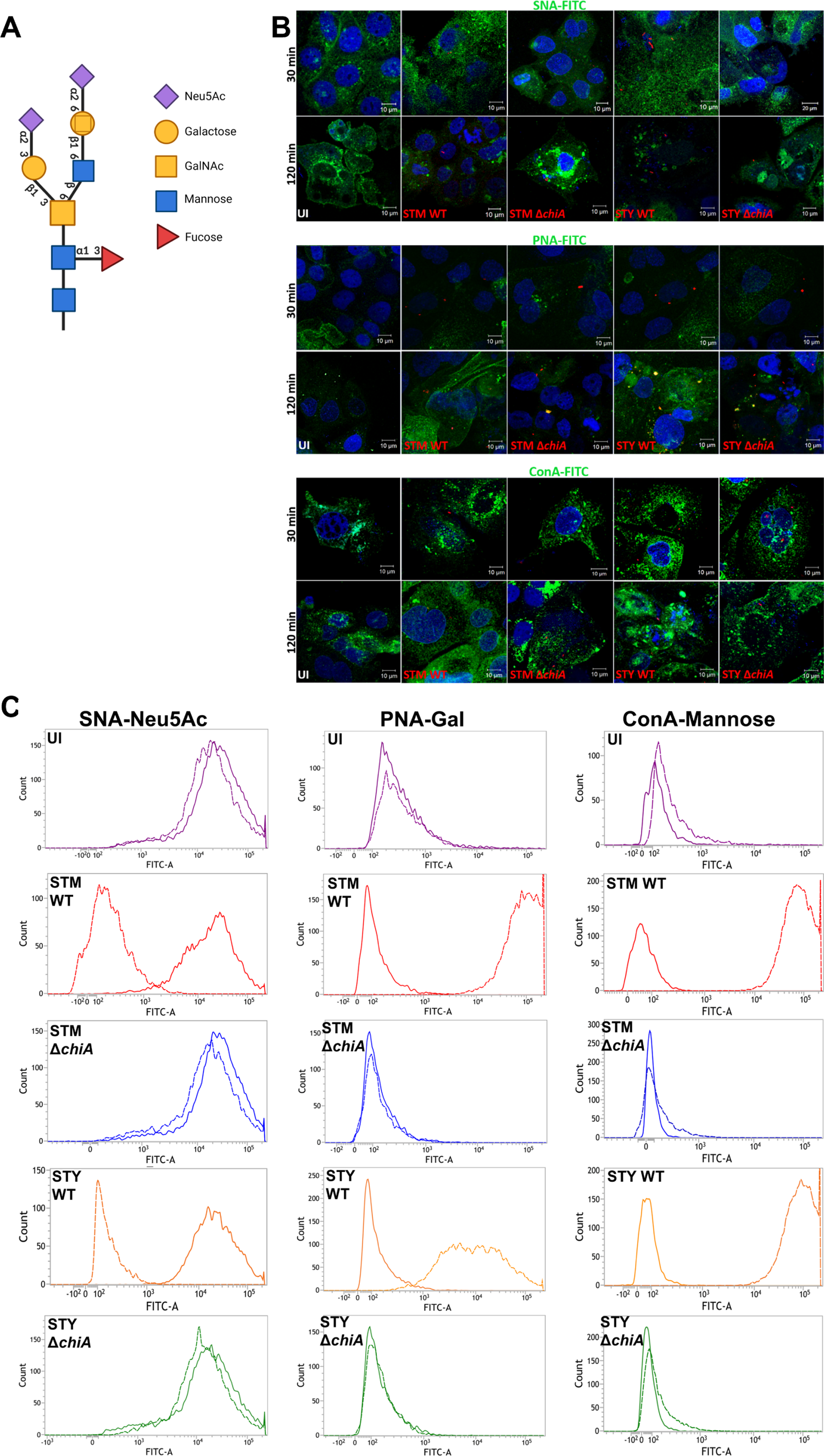
Chitinase helps in glycan remodeling in host epithelial cells. **(A)** Cell surface glycan assembly. **(B)** Representative confocal images of Caco2 cells stained with SNA-FITC (top panel), PNA-FITC (middle panel) and ConA-FITC (bottom panel) lectin after indicated time intervals of STM WT, STM Δ*chiA*, STY WT and STY Δ*chiA* infection (UI-Uninfected). **(C)** Representative flow cytometry histogram showing the cell surface Neu5Ac-bound SNA-FITC (first column), Gal bound PNA-FITC (second column) and mannose bound ConA-FITC (third column) lectin (UI-Uninfected). Solid lines represent MFI after 30 min and dashed lines represent MFI after 120 min. Data are represented as mean ± SEM of 2 independent experiments (N=2).

### *Salmonella* ChiA is required for stabilization of the SCVs in epithelial cells

In the epithelial cells, *Salmonella* resides in a double membrane compartment known as *<u>S</u>almonella*-<u>c</u>ontaining <u>v</u>acuoles (SCVs). *Salmonella* replicates in the epithelial cells by inhibiting the fusion of SCVs with lysosomes. SCVs are specialized endosomes that are marked with the LAMPs, Rab7, Rab11 and vATPases. Since several reports suggested that disruption of SCV leads to bacterial hyperproliferation in the cytoplasm [19], we checked whether the enhanced proliferation of the mutant bacteria after the loss of chitinase is caused by defect in SCV maintenance. We found that after 16 hours of infection, Δ*chiA* mutant bacteria did not co-localize with the late-endosomal marker LAMP1 (**Fig. 3A, S1G, S1H**), suggesting disruption of SCVs in the mutant bacteria infected cells. Upon counting the number of SCV-bound and cytoplasmic bacterial population, we found that 81.6% of STM Δ*chiA* and 87.2% of STY Δ*chiA* quit the vacuole, while very less WT bacteria (STM WT 12.2%, STY WT 8.2%) did not co-localize with LAMP1 (**Fig. 3B-C**). We further quantified the cytosolic bacterial population by chloroquine resistance assay and found significantly higher number of cytosolic mutant bacteria after 16 hours of infection (**Fig. 3D-E**) suggesting that chitinase deletion leads to SCV destabilization in epithelial cells and hyper-proliferation of the cytoplasmic bacteria.

**Fig 3.**
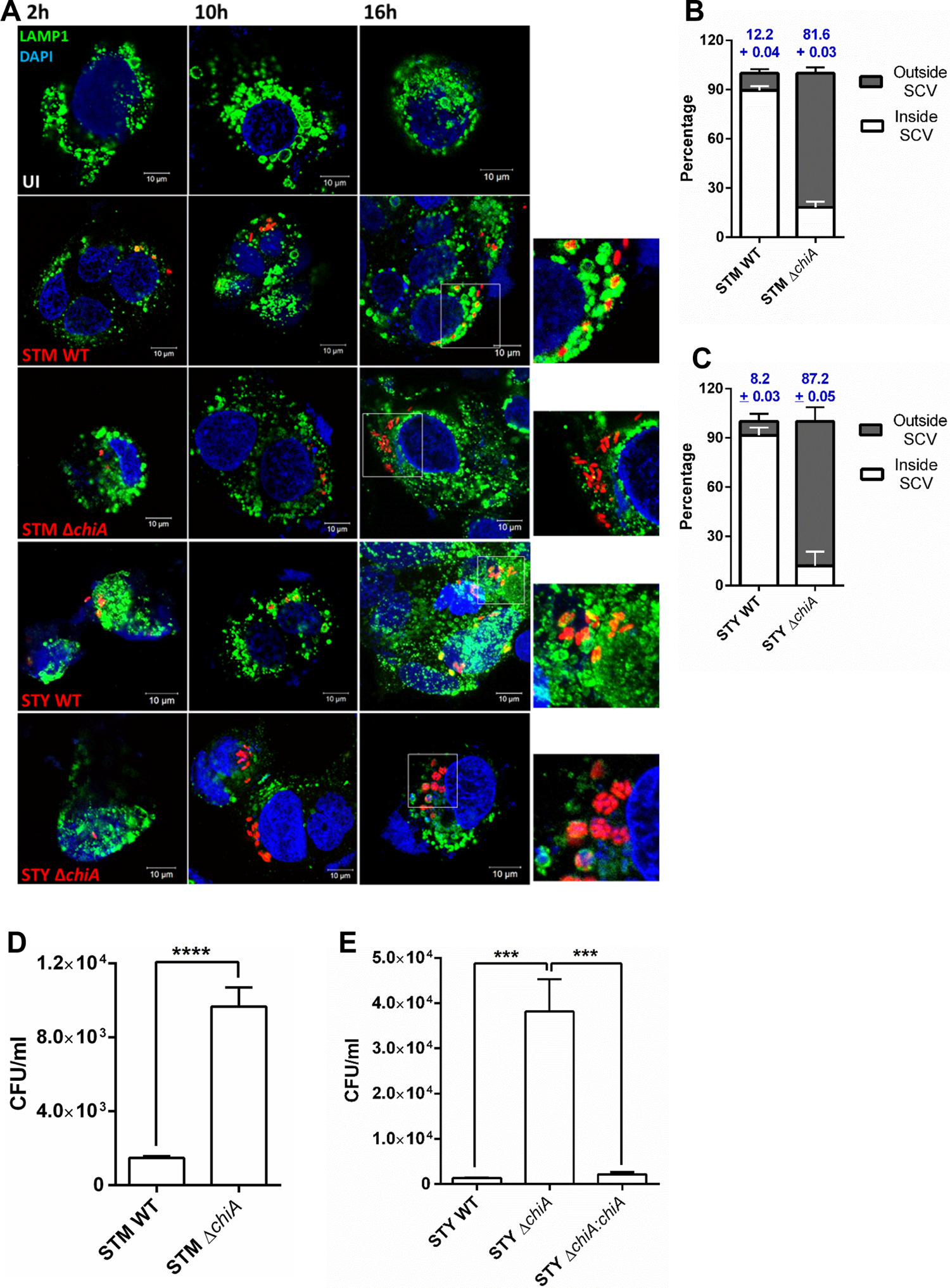
Chitinase deletion destabilizes the *Salmonella*-containing vacuoles in the epithelial cells leading to hyperproliferation of the cytosolic bacteria. **(A)** Representative confocal image of Caco2 cells infected with STM WT, STM Δ*chiA*, STY WT and STY Δ*chiA* strains to visualize the intracellular niche of the bacteria. The SCVs were stained for LAMP1 in presence of 0.01% saponin and 2.5% BSA (UI-Uninfected). **(B)** % of STM WT and STM Δ*chiA,* (C) % of STY WT and STY Δ*chiA* bacteria inside and outside the LAMP1-stained SCVs after 16 hours of infection was calculated. Data are represented as mean ± SEM of 3 independent experiments (N=3). **(D-E)** Absolute CFU/ml values of **(D)** STM WT and STM Δ*chiA*, **(E)** STY WT, STY Δ*chiA* and STY Δ*chiA:chiA* in Caco2 cells in chloroquine resistance assay after 16 hours of infection. Data are represented as mean ± SEM of 3 independent experiments (N=3, n=3). One-way ANOVA was used to analyze the data.

### Chitinase aids in bacterial survival in phagocytes by suppressing antimicrobial responses

After successfully invading the epithelial cells, *Salmonella* reaches lamina propria, where it is taken up by macrophages, DCs and neutrophils which marks the beginning of systemic spread of the pathogen. To understand the role of chitinase in phagocytic cell infection, we infected U937 monocytes and BMDCs. We found that Δ*chiA* mutants were invasion defective in U937 monocytes, while the mutants showed enhanced survival in the monocytes as compared to the WT strains (**Fig. 4A-D, S2A, S2B**). While STM WT and STM Δ*chiA* showed similar invasiveness, surprisingly STY Δ*chiA* showed increased invasion and better survival in BMDCs (**Fig. 4E-H, S2C, S2D**). Higher invasiveness of STY Δ*chiA* in naturally phagocytic cells hints towards that this strain might be highly immunogenic. Phagocytic cells are known to inhibit intracellular bacterial growth by production of reactive nitrogen species and reactive oxygen species [20]. Estimation of nitric oxide produced by the infected BMDCs suggested that Δ*chiA* mutant infected cells produced significantly less nitric oxide (**Fig. 4I**). We further checked the survival of the WT and Δ*chiA* in the absence of NO using NOS2^-/-^ BMDCs and observed that WT bacteria survived similar to the Δ*chiA* mutants (**Fig. 4J-K**), suggesting that chitinase might be involved in the induction of NO in DCs. Furthermore, Δ*chiA* infected peritoneal macrophages (PM) showed significantly less ROS level as compared to WT infected cells (**Fig. 4L**), indicating that chitinase might be regulating RNI and ROS level in the infected cells. NO is an important cell signaling molecule that is produced to kill many human pathogens, such as *Salmonella, Mycobacterium* and *Listeria* etc [20]. Previous studies suggested that low level of NO enhances T cell survival [21], while very high NO is capable of inhibiting T cell proliferation [22]. To check whether lesser production of NO in response to Δ*chiA* mutant infection has any effect on antigen presentation and T cell expansion, we looked into the proliferation of CD8^+^ T cell population using OT1 transgenic mouse (C57BL/6-Tg(TcraTcrb) 1100Mjb/J). The TCR of this transgenic mouse recognizes OVA_257-264_ when presented by MHC-I molecules. This TCR recognition of MHC-I bound cognate peptide results in CD8^+^ T cell proliferation, which can be measured by incorporation of ^3^H thymidine in the DNA of the proliferating population. We found that with Δ*chiA* mutant infection, the expansion of CD8^+^ T cell population was significantly higher in response to the antigen stimulation (**Fig. 4N**). Since APCs such as macrophages and DCs possess MHC-I and MHC-II molecules on the cell surface in order to induce both CD8^+^ T cells and CD4^+^ T cells population, respectively, we detected the surface MHC-II molecules on activated PMs. We found that upon Δ*chiA* infection, the surface MHC-II level was similar to uninfected cells, while WT infection significantly reduced the surface MHC-II level (**Fig. 4O, S2E**). Further immunofluorescence analysis verified that Δ*chiA* infection did not change the surface MHC-II molecules on macrophages (**Fig. 4M, S2F**). Together these data suggests that *Salmonella* ChiA dampens host antimicrobial responses leading to enhanced pathogen survival.

**Fig 4.**
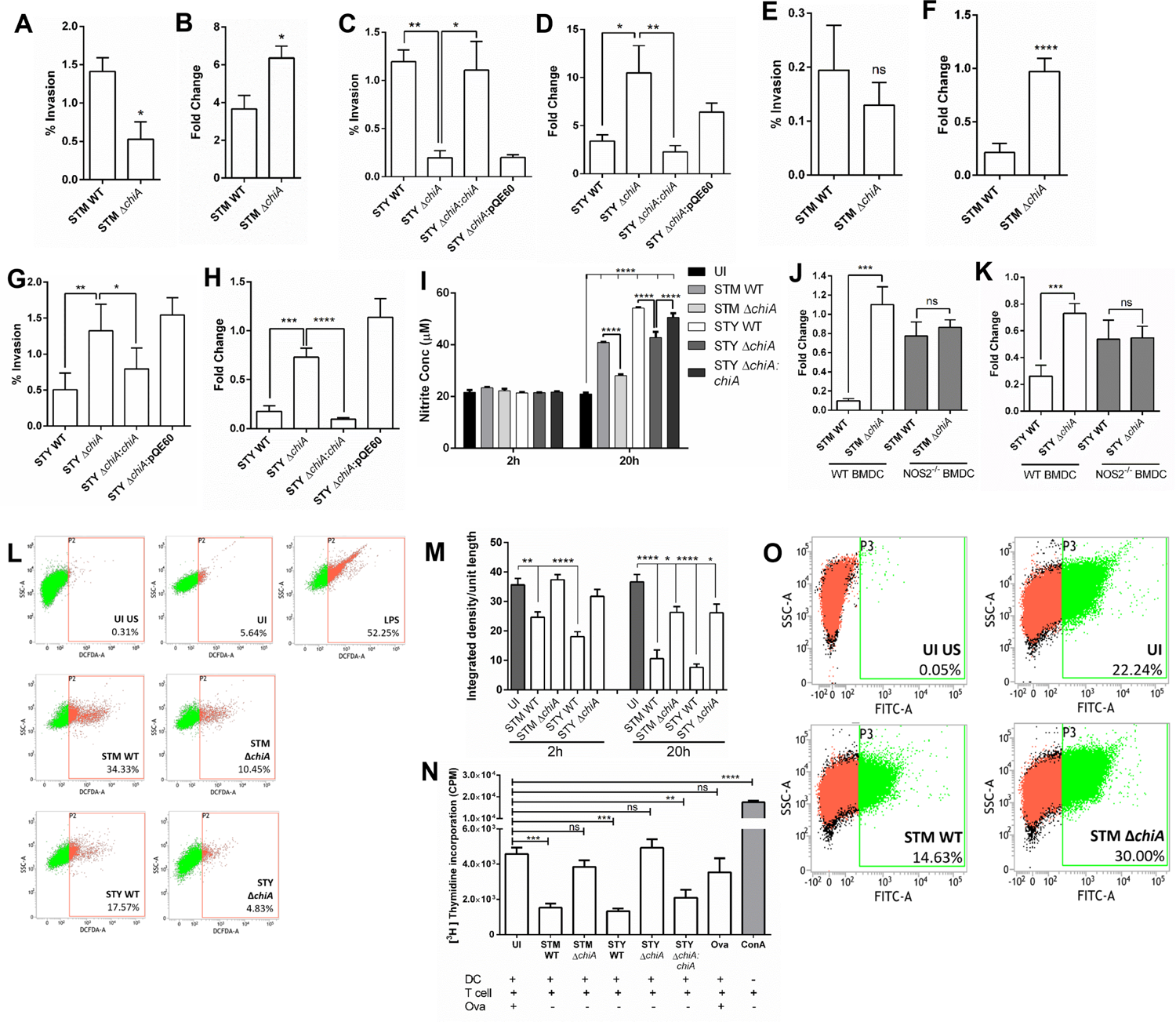
Chitinase induces NOS and ROS generation in phagocytic cells and inhibits of antigen presentation. **(A)** % invasion and **(B)** fold proliferation of STM WT and STM Δ*chiA* strains in U937 derived monocytes by gentamicin protection assay. Data are represented as mean ± SEM of 3 independent experiments (N=3, n=3). Unpaired Student’s t test was used to analyze the data. **(C)** % invasion and **(D)** fold proliferation of STY WT, STY Δ*chiA,* STY Δ*chiA:chiA* and STY Δ*chiA:*pQE60 strains in U937 derived monocytes by gentamicin protection assay. Data are represented as mean ± SEM of 3 independent experiments (N=3, n=3). One-way ANOVA was used to analyze the data. **(E)** % invasion and **(F)** fold proliferation of STM WT and STM Δ*chiA* strains in U937 derived monocytes by gentamicin protection assay. Data are represented as mean ± SEM of 3 independent experiments (N=3, n=3). Unpaired Student’s t test was used to analyze the data. **(G)** % invasion and **(H)** fold proliferation of STY WT, STY Δ*chiA,* STY Δ*chiA:chiA* and STY Δ*chiA:*pQE60 strains in U937 derived monocytes by gentamicin protection assay. Data are represented as mean ± SEM of 3 independent experiments (N=3, n=3). One-way ANOVA was used to analyze the data. **(I)** Extracellular NO was estimated by Greiss assay from conditioned media obtained from STM WT, STM Δ*chiA*, STY WT, STY Δ*chiA* and STY Δ*chiA:chiA* infected DCs. (UI-Uninfected). Data are represented as mean ± SEM of 3 independent experiments (N=3, n=3). Two-way ANOVA was used to analyze the data. **(J-K)** Intracellular survival of **(J)** STM WT and STM Δ*chiA* and **(K)** STY WT and STY Δ*chiA* strains were calculated in WT and NOS2^-/-^ BMDCs by gentamicin protection assay. Data are represented as mean ± SEM of 3 independent experiments (N=3, n=3). Unpaired Student’s t test was used to analyze the data. **(L)** Representative flow cytometry plot for ROS estimation by DCFDA assay from STM WT, STM Δ*chiA*, STY WT, STY Δ*chiA* and STY Δ*chiA:chiA* infected and LPS treated peritoneal macrophages. (UI US-Unstained uninfected, UI-Uninfected. LPS-Lipopolysaccharide). **(M)** Quantification of the MHC-II density per unit length of the cell membrane of STM WT, STM Δ*chiA*, STY WT, STY Δ*chiA* and STY Δ*chiA:chiA* infected PMs after indicated time (UI-Uninfected). Data are represented as mean ± SEM of 2 independent experiments. One-way ANOVA was used to analyze the data. (UI US-Unstained uninfected, UI-Uninfected). **(N)** ^3^H thymidine incorporation assay to assess CD8^+^ T cell proliferation after 20 hours of infection with STM WT, STM Δ*chiA*, STY WT, STY Δ*chiA* and STY Δ*chiA:chiA* (UI-Uninfected, OVA-Ovalbumin, ConA-Concanavalin A). Data are represented as mean ± SEM of 3 independent experiments. One-way ANOVA was used to analyze the data. **(O)** Representative flow cytometry plot showing the level of surface MHC-II molecules on PMs infected with STM WT and STM Δ*chiA* for 20 hours (UI US-Unstained uninfected, UI-Uninfected).

### Chitinase facilitates *in vivo* invasion, survival and pathogenesis of *Salmonella* Typhimurium

To further delineate the role of chtitinase in *Salmonella* infection *in vivo*, we orally infected 6 weeks old C57BL/6J mice with lethal dose of the bacterial strains (10^8^ CFU/animal) and observed for survival. *Salmonella enterica* serovar Typhimurium mimics the systemic typhoidal disease in murine model. STM Δ*chiA* infected cohort showed enhanced animal survival as compared to the STM WT infected cohort (**Fig. 5A**), suggesting a role of chitinase A during infection *in vivo*. We also found that STM Δ*chiA* mutant bacteria were shed prior to the STM WT and the Δ*chiA* mutant was defective in PP colonization after 2 hours of oral gavage (**Fig. 5B-C**). Further, we orally infected C57BL/6J mice with sublethal dose of *Salmonella* strains (10^7^ CFU/animal) and bacterial CFU from liver, spleen, mesenteric lymph node (MLN) and PP was enumerated after sacrificing the animals after indicated time intervals. We found that STM Δ*chiA* mutant infected animals showed less bacterial burden in each of the organs and higher body weight as compared to the STM WT infected animals (**Fig. 5D-H**). Since our *ex vivo* data suggested that the Δ*chiA* mutant was unable to induce NO production which is known to affect T cell survival, we checked whether chitinase mediated NO upregulation has any effect *in vivo*. We also found a significant increase in the spleen length after 20 days of infection with STM Δ*chiA* bacteria, as compared to the STM WT infected mice (**Fig. S3A, S3B**). We isolated total splenocytes from these infected spleens and checked the activated CD4^+^ T cell population and T cell proliferation by flow cytometry. We found that STM Δ*chiA* infection leads to a significant increase in CD4^+^ T cell population, as well as an increase in the activated CD25^+^ T cells (**Fig. 5K**). Analysis of T cell mediated cytokine response revealed that there was a significant increase in the pro-inflammatory cytokines IL2 and IFNγ, in the serum isolated from Δ*chiA* infected animals (**Fig. 5I-J**), whereas there was no difference in the anti-inflammatory cytokine levels (**Fig. S3C, S3D**). Lesser IL2 in STM WT infected mice serum as compared to that in Δ*chiA* mutant infected serum further strengthens the previous finding that ChiA aids in dampening of T cell activation, as IL2 is a key marker of CD8^+^ T cell proliferation [23]. Previous reports suggested that high level of IFNγ can induce B cell proliferation and enhance IgG2a and IgG3 production [24]. Therefore, we looked into the role of chitinase in anti-*Salmonella* immune response by detecting the anti-*Salmonella* IgG titer from infected mice serum. Interestingly, we found a significant increase in the anti-*Salmonella* antibody titer in the serum obtained from STM Δ*chiA* mutant infected cohort (**Fig. 5L**). We further used the polyclonal convalescent sera isolated from STM Δ*chiA* infected mice to probe against STM WT-mCherry whole cell lysate to test the polyclonality of the sera. Multiple dense bands against various *Salmonella* proteins were obtained after incubating the membrane with sera collected from STM Δ*chiA* mutant infected cohort (**Fig. S3E**). Together these data suggest that *Salmonella* chitinase A is essential for restricting innate and humoral immune responses against *Salmonella*.

**Fig 5.**
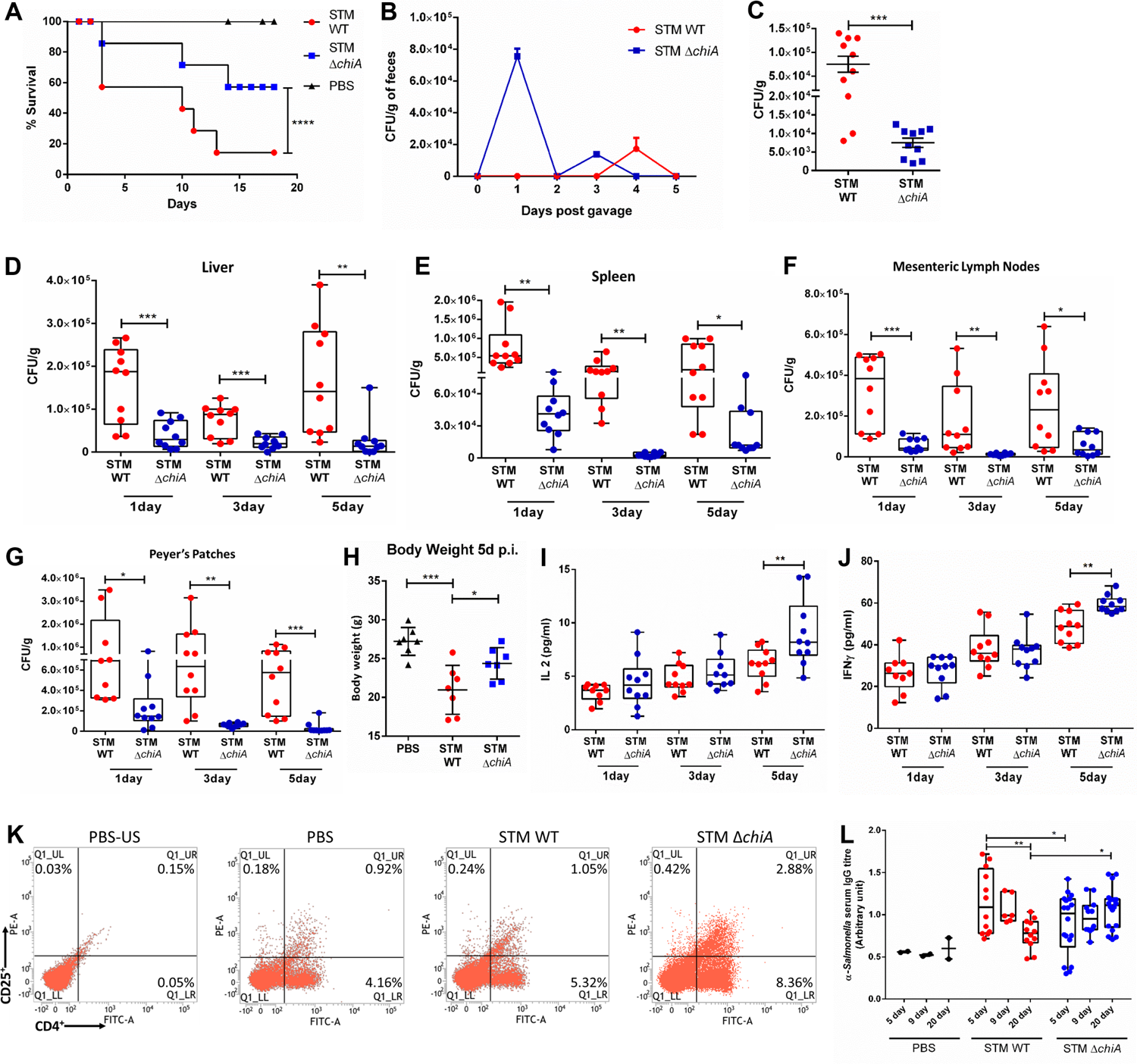
Chitinase facilitates *in vivo* invasion, survival and pathogenesis of *Salmonella* Typhimurium. **(A)** Survival of the mice infected with lethal dose of STM WT and STM Δ*chiA* (PBS= Phosphate Buffered Saline). Data are presented from one independent experiment, representative of 3 independent experiments (N=3). **(B)** Bacterial shedding in the feces of the animals infected with STM WT and STM Δ*chiA*. Data are presented as mean ± SD of one independent experiment, representative of 3 independent experiments (N=3). **(C)** *In vivo* invasion in Peyer’s patches 2 hour post oral gavage by either STM WT or STM Δ*chiA*. Data are presented as mean ± SEM of 3 independent experiments. Unpaired Student’s t test was used to analyse the data. **(D-G)** Bacterial burden in **(D)** liver, **(E)** spleen, **(F)** MLN and **(G)** PP of the infected mice on different days post infection with sublethal dose of STM WT and STM Δ*chiA*. Data are presented as mean ± SEM of 3 independent experiments. Unpaired Student’s t test was used to analyze the data. **(H)** Body weight of the infected mice was measured 5 days post infection with sublethal dose of STM WT and STM Δ*chiA*. Data are presented from 3 independent experiments. (PBS-Phosphate Buffered Saline). **(I-J)** Pro-inflammatory cytokines **(I)** IL2 and **(J)** IFNγ level in serum from STM WT and STM Δ*chiA* infected mice after indicated time. Data are presented as mean ± SEM of 3 independent experiments (N=3). One-way ANOVA was used to analyze the data. **(K)** Flow cytometry analysis of total splenocytes isolated from the spleens isolated from STM WT and STM Δ*chiA* infected mice after 20 days. Splenocytes were stained for CD4 and CD25 markers (US-PBS-Unstained splenocytes from PBS infected mouse). Data are presented from one independent experiment, representative of 3 independent experiments (N=3). **(L)** Serum anti-*Salmonella* antibody level was checked by sandwich ELISA after indicated time. Data are presented as mean ± SEM of 3 independent experiments (N=3). Two-way ANOVA was used to analyze the data.

### Chitinase helps in *Salmonella* Typhi pathogenesis in *C. elegans*

*Salmonella* Typhi is a human obligatory pathogen that does not cause a significant infection in mice because of the presence of TLR11 [25]. Long before TLR11^-/-^ mice model came into existence, Labrousse *et al.* suggested *Caenorhabditis elegans* can be used as an alternative host to study *S.* Typhi pathogenesis [26]. Given that *C. elegans* pharyngeal lumen is rich in chitin, it served as a suitable host to study the role of chitinase in bacterial pathogenesis [27]. We began with checking the bacterial CFU in the infected worms after 24 hours and 48 hours of continuous feeding. We found that the STY Δ*chiA* and STY Δ*chiA:*pQE60 strains showed a higher bacterial burden after 24 hours of continuous feeding (**Fig. S3F**), but the fold change of bacteria were lesser than that of the STY WT and STY Δ*chiA:chiA* strains (**Fig. 6A**). We further checked animal survival after infecting the worms with different bacterial strains and found that all the *Salmonella* Typhi strains are pathogenic to the animals as compared to *E. coli* OP50, while STY Δ*chiA* showed slower death (TD_50_ 330±8hrs) in the worms as compared to the STY WT (TD_50_ 190±10hrs) and STY Δ*chiA:chiA* (TD_50_ 270±12hrs) strains (**Fig. 6B**). Together these data suggest that chitinase deletion renders *Salmonella* Typhi less virulent in *C. elegans* infection. We further checked bacterial colonization in the worm’s gut using the transgenic worm FT63 strain that expresses GFP in the epithelial cells. We visualized the bacterial colonization in the worms gut after 24 hours and 48 hours of continuous feeding. We used STM Δ*invC* mutant as a control which is known to be invasion defective in nonphagocytic cells [28]. We found that *S.* Typhi Δ*chiA* showed less colonization than STY WT after 24 hours continuous feeding, while the colonization was significantly reduced after 48 hours feeding (**Fig. 6C**), suggesting chitinase is required for successful gut colonization in *C. elegans*. Percent colonization was measured as the ratio of the diameter of the lumen occupied by the bacteria to the total diameter of the gut (**Fig. S3G**). Interestingly STM Δ*invC* did not show any defect in colonization in the *C. elegans* gut, suggesting SPI1 effector InvC is not essential for colonization in the worms gut. We next checked if *S.* Typhi utilizes chitinase to colonize the chitin-rich pharyngeal lumen by infecting N2 worms with different strains of *Salmonella* and stained the chitin-rich parts of the worms using eosin Y. We found that after 24 hours of continuous feeding, luminal STY WT and STM Δ*invC* bacteria colocalized with the chitin-rich regions of the pharyngeal wall and terminal bulb (grinder), whereas STY Δ*chiA* bacteria did not show any colocalization with the pharyngeal wall (**Fig. 6D**), suggesting *Salmonella* Typhi utilizes chitinase to colonize the chitin-rich pharynx and terminal bulb. Additionally, we looked into the role of chitinase in bacterial persistence in the worms gut by infecting the worms with different STY strains, followed by feeding onto *E. coli* OP50. We hypothesized that if a bacterial strain can adhere to the gut lumen effectively, it will remain adhered to the gut lumen and proliferate even when the worms are fed with *E. coli* OP50. We observed that after 24 hours of feeding on STY Δ*chiA* followed by 24 hours feeding on *E. coli* OP50, the STY Δ*chiA* was unable to persist in the gut, whereas STY WT showed significantly higher colonization in the pharyngeal lumen (**Fig. S3H**). When we further extended the infection for 48 hours, followed by 24 hours of *E. coli* OP50 feeding, we observed that STY WT showed profound colonization of the gut lumen, while STY Δ*chiA* colonization was diminished (**Fig. 6E**), suggesting *Salmonella* utilizes chitinase to attach to the lumen wall for enhanced persistence in the worms. Interestingly, after 24 hours of continuous feeding STY WT attached to the luminal wall, but not STY Δ*chiA* strain. (**Fig. S4A**). After 48 hours of continuous feeding, we detected STY WT and STM Δ*invC* bacteria in the extra-intestinal tissues of the worms, while STY Δ*chiA* did not show extra-intestinal colonization (**Fig. 7A, S4A, S4B**), suggesting that chitinase might be required to invade extra-intestinal tissues of the worms. To the best of our knowledge, this study is the first report suggesting an extra-intestinal invasion/colonization by *Salmonella* Typhi in *C. elegans*.

**Fig 6.**
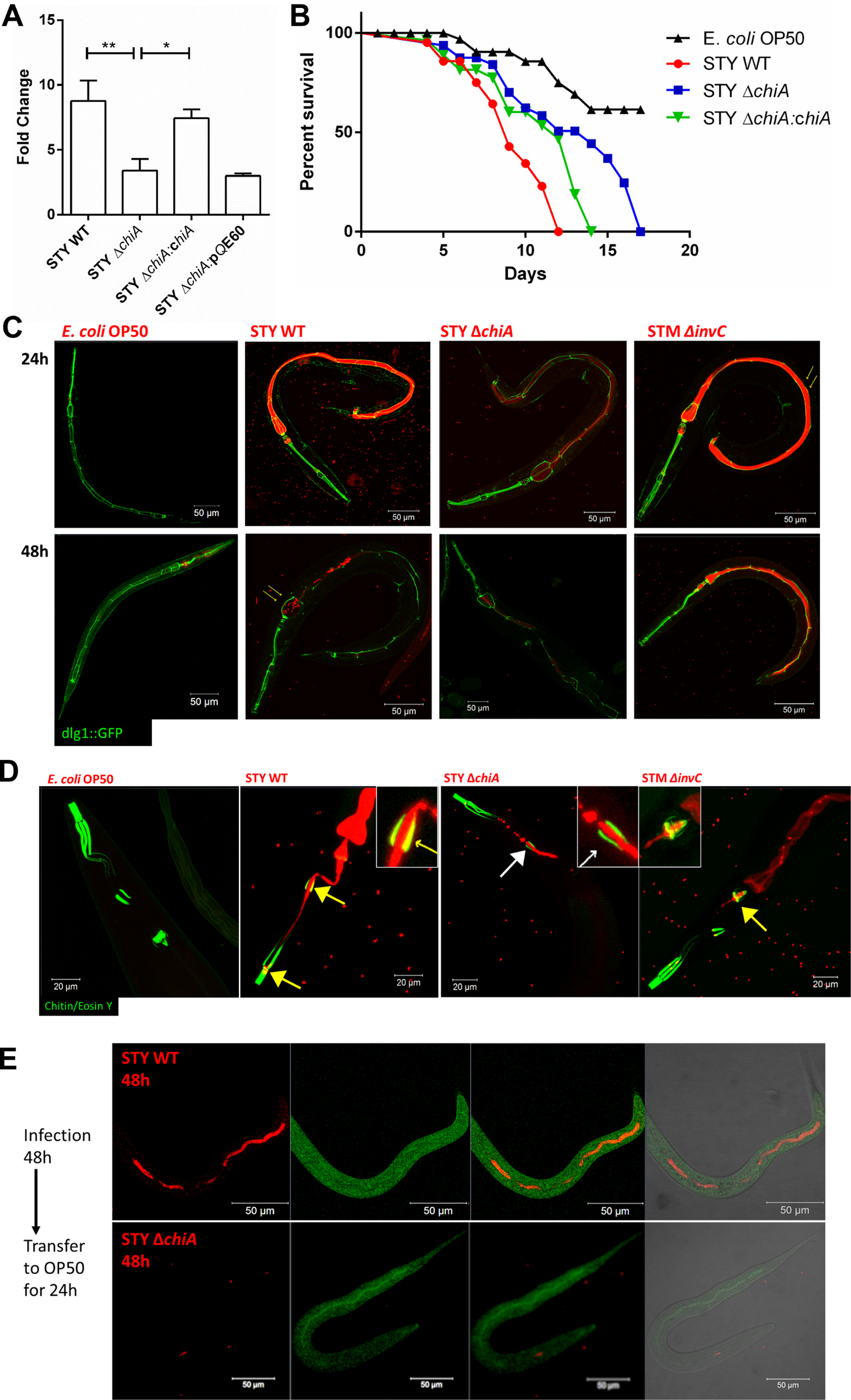
Chitinase enhances *Salmonella* Typhi pathogenesis in *C. elegans.* **(A)** Bacterial proliferation measured by the ratio of bacterial CFU obtained from infected *C. elegans* after 24 hours and 48 hours continuous feeding on STY WT, STY Δ*chiA*, STY Δ*chiA:chiA* and STY Δ*chiA*:pQE60 strains. Data are represented as mean ± SEM of 4 independent experiments. One way ANOVA was used to analyze the data. **(B)** Survival of the worms fed on *E. coli* OP50, STY WT, STY Δ*chiA* and STY Δ*chiA:chiA*. Data are presented from one independent experiment, representative of 4 independent experiments. **(C)** Representative confocal images of bacterial colonization in worms gut as observed by infecting transgenic FT63 worms by bacteria expressing mCherry for indicated time. Yellow arrows show presence of intact bacteria in the terminal bulb of the worm. **(D)** Representative confocal images of bacterial colonization on the chitin-rich organs of the worms as detected by eosin Y staining. Yellow arrows colocalization of the bacteria (red) and the eosin-stained chitin containing regions (green). White arrow shows absence of colocalization of the bacteria and chitin-rich organs. **(E)** Representative confocal images showing bacterial colonization in the worms gut after 48 hours of STY WT and STY Δ*chiA* feeding followed by feeding on *E. coli* OP50 for subsequent 24 hours.

**Fig 7.**
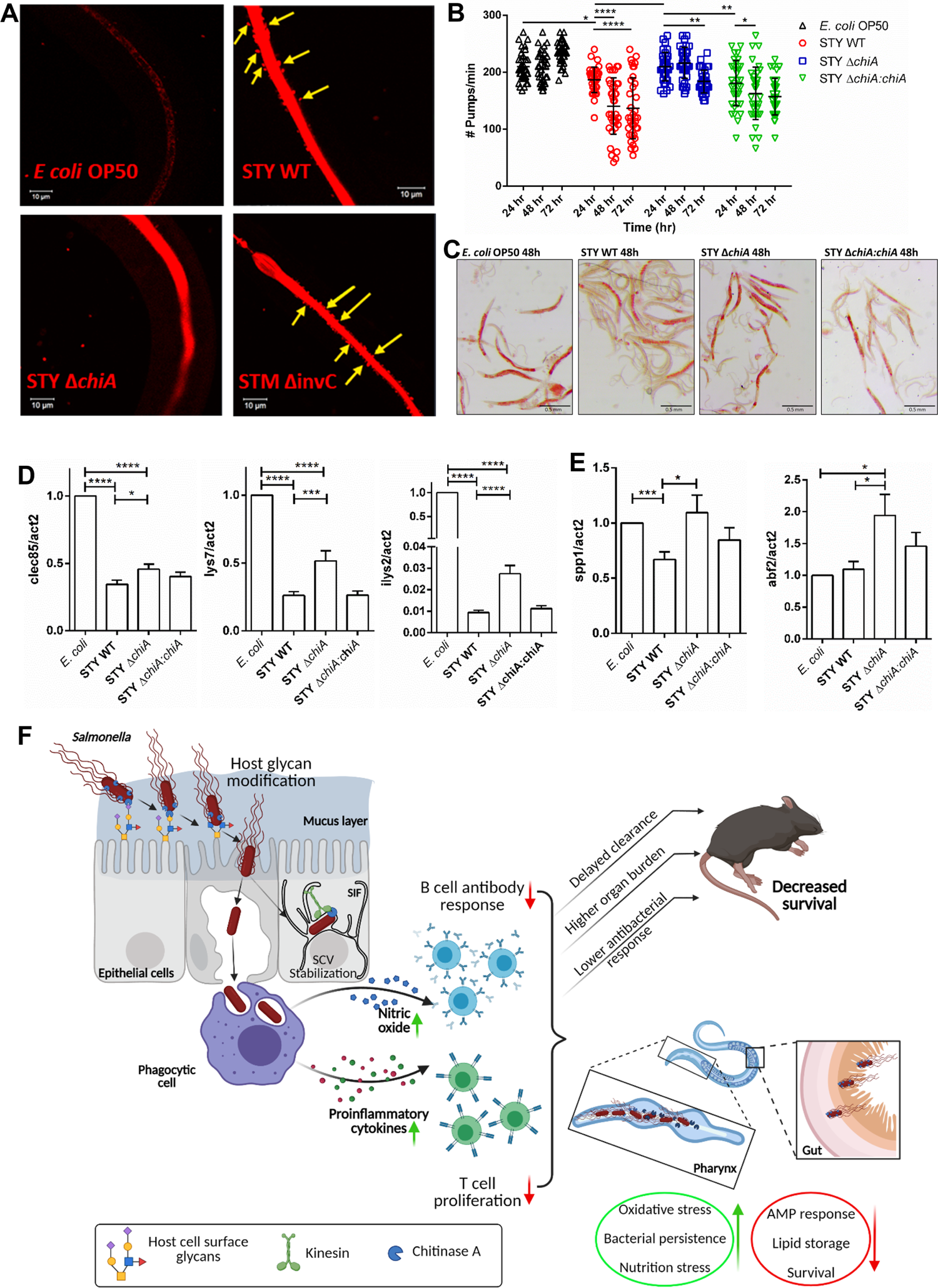
*Salmonella* chitinase is important for alteration of metabolism and antibacterial defense in *C. elegans.* **(A)** Representative images of bacterial colonization of the worms gut at higher magnification. Yellow arrows show presence of the STY WT and STM Δ*invC* bacteria outside the gut lumen. **(B)** Quantification of no. of pharyngeal pumps/min in *E. coli* OP50, STY WT, STY Δ*chiA* and STY Δ*chiA:chiA* fed worms. Data are represented as mean ± SD of 3 independent experiments. Two-way ANOVA was used to analyze the data. **(C)** Representative images of Oil Red O stained worms fed with *E. coli* OP50, STY WT, STY Δ*chiA* and STY Δ*chiA:chiA* for 48 hours. **(D-E)** Quantitative RTPCR analysis of the p38 MAPK dependent antimicrobial peptide genes **(D)** *clec85, lys7* and *lys2* **(E)** *spp1* and *abf2* in worms fed with *E. coli* OP50, STY WT, STY Δ*chiA* and STY Δ*chiA:chiA* for 48 hours. Fold change was normalized over *act2.* Data represents mean ± SEM of 4 independent experiments. One-way ANOVA was used to analyze the data. **(F)** Model depicting the role of Chitinase A in bacterial invasion and regulation of host immune response during *Salmonella* pathogenesis in mouse and *C. elegans* host (Created with Biorender.com).

### *Salmonella* chitinase is important for alteration of metabolism and antibacterial defense in *C. elegans*

Grinder, a part of the terminal bulb, is the complex structure that helps in uptake and grinding of bacteria before it passes to the intestine where the nutrients get absorbed. In a healthy and well-fed state, worms feed at the average rate of 200 pumps/min. Since we observed that *Salmonella* uses chitinase to colonize chitin-rich organs (**Fig. 6D**), we next looked into the nutritional state of the worms by counting the number of pharyngeal pumps per min. We found a significant reduction in the number of pharyngeal pumps/min after 72 hours of STY WT and STY Δ*chiA:chiA* infection (**Fig. 7B**). Further, *in vivo* oxidative stress was quantified using CL2166 worms, that possess oxidative stress inducible GFP. STY WT and STY Δ*chiA:chiA* infected worms showed significantly higher oxidative stress and ‘bag of worms’ phenotype (**Fig. S4C, S4D**). We next checked the nutritional fitness of the worms by measuring the lipid content using oil red O staining. Oil red O (ORO) is a fat-soluble dye that stains neutral lipids [29]. Interestingly we observed significant fat-loss in the worms fed with STY WT and STY Δ*chiA:chiA* for 24 hours and 48 hours as compared to STY Δ*chiA* infected wormed (**Fig. 7C, S4E**). We further checked whether chitinase deletion has any effect on the immune response of the worms. Although our data suggest both STY WT and STY Δ*chiA* downregulated p38 MAP kinase pathway genes *pmk1* and *mek1* equally (**Fig. S4F**), p38 MAP kinase pathway regulated antimicrobial peptides were differentially expressed. While *clec85, lys7, ilys2* expressions were severely downregulated in both STY WT and STY Δ*chiA* infection, all of them showed significant rescue in the worms infected with STY Δ*chiA* bacteria (**Fig. 7D**). Interestingly, we observed that STY WT infection induced downregulation of antimicrobial peptide *spp1* was completely rescued upon STY Δ*chiA* infection, while *abf2* was significantly upregulated upon infection with STY Δ*chiA* (**Fig. 7E**), indicating an important contribution of chitinase in dampening the antimicrobial responses of the host.

## Discussion

*Salmonella* is a facultative intracellular human pathogen that has co-evolved with its host and has also developed various strategies to evade the host’s immune responses. The detailed understanding of the metabolism and the ease of genetic manipulation has made *Salmonella* an excellent the model to study the role of metabolism related proteins in the light of host-pathogen interaction. Although *Salmonella* pathogenesis is governed by classical virulence factors such as adhesins, invasins and toxins, emerging reports suggest that various unique metabolic proteins are important in various aspects of *Salmonella* pathogenesis. Several reports suggest that *Salmonella* can utilize a large pool of chemically diverse host nutrients, such as carbohydrates, lipids, amino acids etc [30]. One such carbon metabolism related protein encoding gene *chiA* (STM0022) was found to be highly upregulated in intracellular *Salmonella* Typhimurium str. SL1344 isolated from infected macrophages and epithelial cells [11, 12]. Bacterial chitinases belong to GH18 and GH19, which are getting recognized as bacterial virulence factors along with several other structurally similar glycosidases such as sialidases, muraminidases, N-acetylgalactosidases etc [2]. Although *Salmonella chiA* was upregulated during infection, the role of this chitinase in *Salmonella* pathogenesis remains elusive. To answer this question, we generated isogenic Δ*chiA* mutant by one-step gene inactivation method. We did not find any significant difference in the *in vitro* growth among the two strains. To test whether *chiA* upregulation in infected cells has any significance in the pathogenesis, we infected epithelial cells and phagocytes with the mutant strain. Interestingly, we found that the mutant was invasion defective in epithelial cells. Previous reports suggested that *Salmonella* remodels the host cell surface glycans to facilitate invasion in the epithelial cells [13–15]. Our observations from the lectin-binding assay suggests that chitinase aids in glycan remodeling by cleaving the terminal glycosyl molecules and making the mannose residues accessible to the bacteria for binding. We further found that absence of ChiA leads to destabilization of the SCV and hyper-proliferation of the mutant bacteria in the cytoplasm of the epithelial cells. The Δ*chiA* mutants survived better than WT strains in the phagocytes, suggesting the Δ*chiA* mutants were protected from phagocytes mediated bacterial killing. Additionally, we found that the phagocytes infected with the mutant bacteria produced less antimicrobial molecules such as NO and ROS. We also found that ChiA was important for downregulating the MHC-I molecules on the dendritic cells, leading to the inhibition of CD8^+^ T cell proliferation and subsequent antigen presentation. In coherence with the available literatures [22], the enhanced T cell proliferation could be attributed to the absence of NO induction by the Δ*chiA* mutant strains. We further showed that absence of *chiA* failed to downregulate the surface MHC-II molecules on the activated macrophages, which is a well-known phenomenon during *Salmonella* infection [31]. *In vivo* infection in C57BL/6 mice suggested that STM Δ*chiA* mutant was unable to invade the Peyer’s patches, leading to an early fecal shedding and enhanced pathogen clearance. STM Δ*chiA* mutant infected cohort presented significantly less bacterial burden in the liver, spleen, MLN and PP, as well as they showed increased survival, suggesting that the STM Δ*chiA* mutant is less virulent *in vivo.* Analysis of total splenic lymphocytes by flow cytometry suggested that the Δ*chiA* mutant infected cohort had an increased activated T cell population (CD4^+^CD25^+^) in the spleens, suggesting an intensified immune response in these mice. This was corroborated by significant increment in the pro-inflammatory cytokines and anti-STM IgG antibody titer in the STM Δ*chiA* infected mice sera. In the invertebrate *C. elegans* model, chitinase helps in bacterial attachment to the pharyngeal lumen. Additionally, we found that the chitinase helps in *Salmonella* Typhi colonization and persistence in the worms, since deletion of *chiA* leads to bacterial clearance from the worm’s gut. In addition, our data suggest that *Salmonella* Typhi chitinase might help in extra-intestinal tissue invasion in the worms. Chitinase was found to be regulating the fat-responsive immune response such as antibacterial peptide synthesis in the worms. We found significantly higher expression of antimicrobial peptides genes *spp1* and *abf2* when the worms were infected with STY Δ*chiA* strain, hinting towards a potential role of chitinase in modulating innate immune response in the worms. Together our data suggest that *Salmonella* Chitinase regulates different aspects of pathogenesis, ranging from aiding in invasion in the epithelial cells, impairing the activity of professional antigen presenting cells to as diverse as immune response regulation in various hosts (**Fig. 7F**) and emerges as a novel virulence factor.

## Materials and Methods

### Bacterial strains

All *Salmonella* Typhimurium strains used in this study are listed below with their genetic description. *Salmonella enterica* serovar Typhimurium strain 14028S was used as the wild type strain, and was also the parental background for all the mutant strains used in this study, i.e. Δ*chiA* and Δ*invC*. All strains were grown and maintained in Lennox broth (LB; 0.5% NaCl, 1% casein enzyme hydrolysate and 0.5% yeast extract) at 37°C under shaking conditions. *Salmonella enterica* serovar Typhi strain CT18 was used as the wild type strain, and was also the parental background for the mutant strain used in this study, i.e. Δ*chiA*. *S*. Typhi *chiA* was trans-complemented in pQE60 plasmid in 5’ NotI-*chiA*-BamHI 3’ direction. This plasmid was transferred to STY Δ*chiA* strain to make complement strain. Complemented strain STY Δ*chiA:chiA* and empty vector strain STY Δ*chiA;*pQE60 strains were maintained on LBA supplemented with ampicillin (50 µg/ml). The mCherry expressing strains were cultured in Lennox broth with 50µg/ml Ampicillin at 37°C in shaking condition.

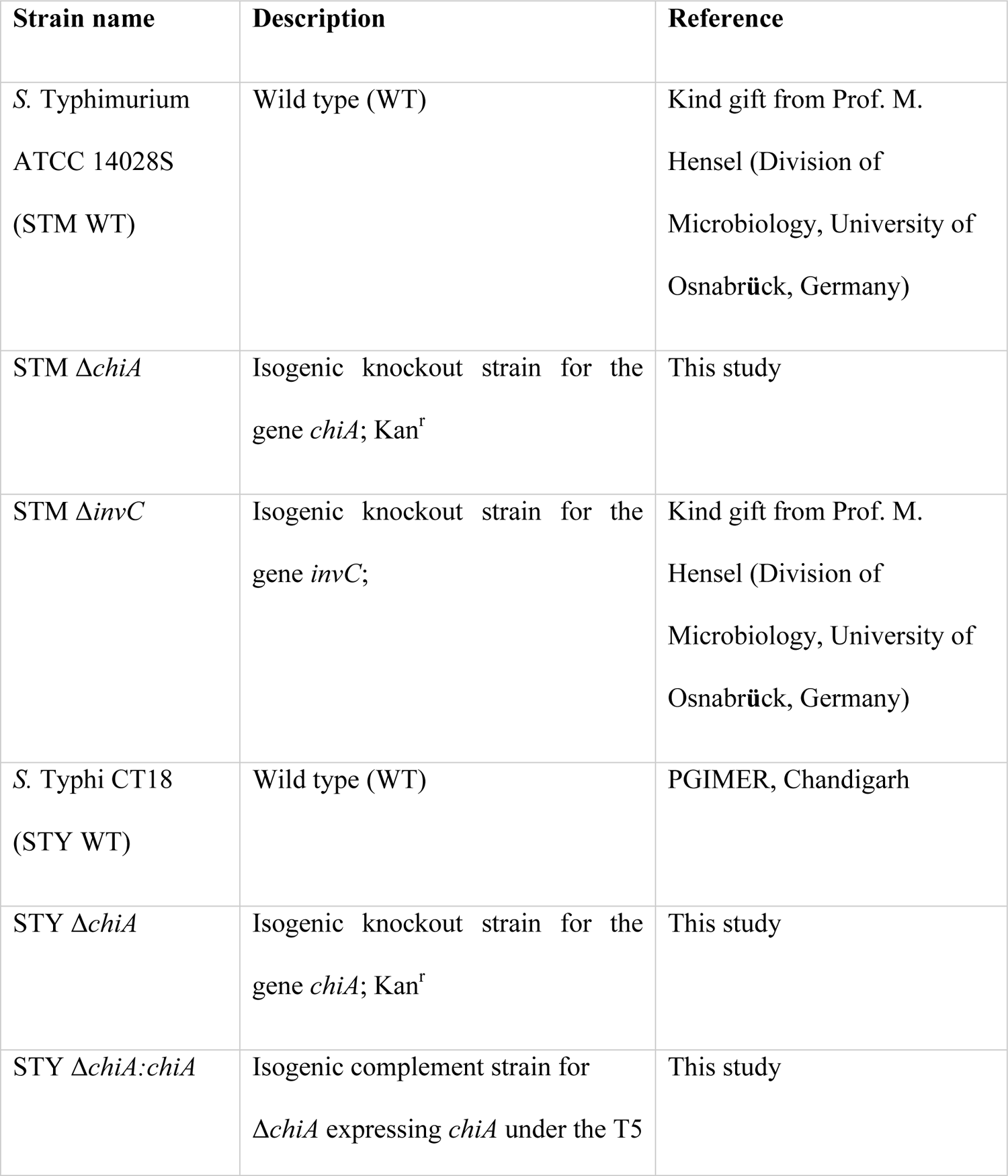
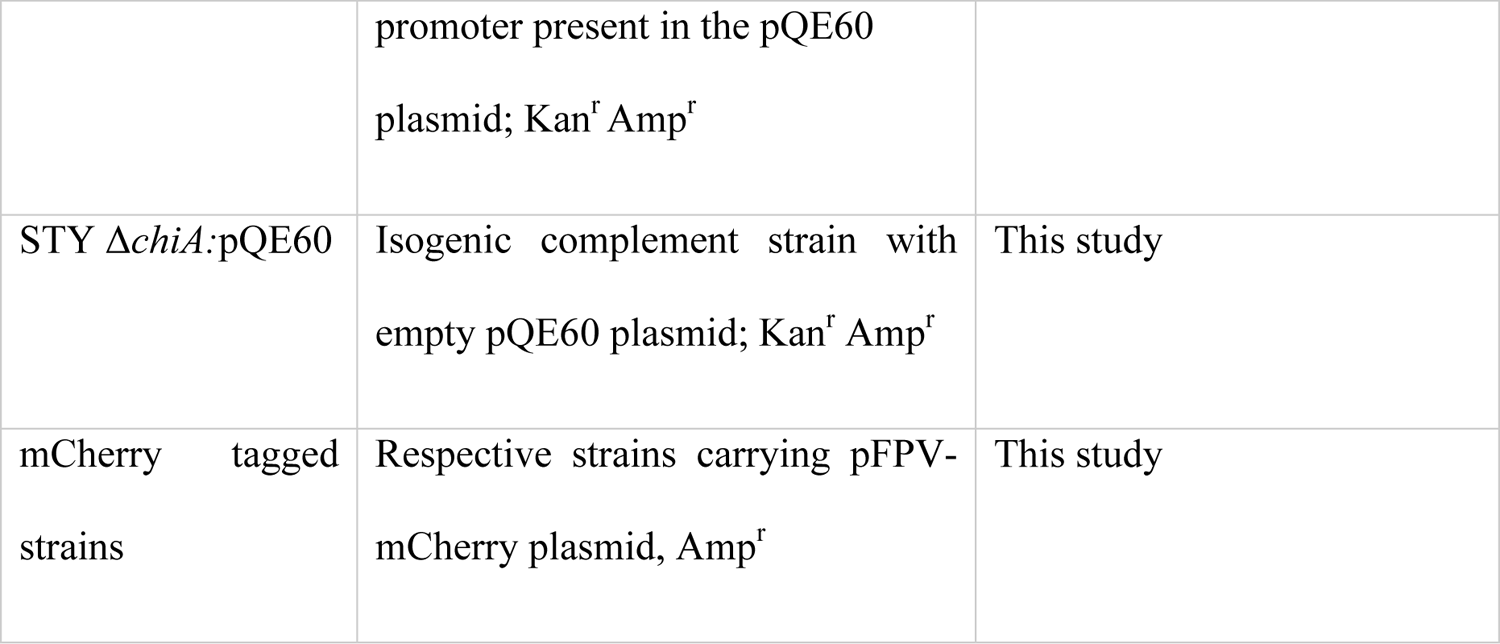
List of strains used in this study.

### Isolation and maintenance of primary cells and cell lines

Human colorectal adenocarcinoma cell line Caco2 (ATCC HTB-37) was cultured in complete DMEM media (Lonza), whereas human monocyte cell line U937 (ATCC CRL-1593.2) was maintained in complete RPMI 1640 media (Lonza) with 100 µM β-mercaptoethanol and differentiated to macrophages using 20 ng/ml PMA for 24 hours prior to infection. Bone-marrow was isolated from either wildtype (NOS2^+/+^) C57BL/6J mice or NOS2^-/-^ C57BL/6J mice as described previously [31]. Briefly, tibia and femur bones were carefully taken out, caps were removed and the marrow was flushed with RPMI 1640 media using a 26G needle. After making single cell suspension, RBCs were lysed using RBC lysis buffer. Cells were pelleted and grown in complete RPMI 1640 media supplemented with 20 ng/ml mGM-CSF (Peprotech), antibiotics and 100 µM β-mercaptoethanol. After every 2 days, the media was replenished. Once approximately 65-70% of the cells were differentiated to dendritic cells (loosely adherent spheres), the cells were collected and used for further experiments. To obtain peritoneal macrophages (PMs), thioglycolate was injected in the peritoneal cavity of C57BL/6J mice. After 5 days these mice were sacrificed and ice cold PBS was injected in the peritoneum to collect the peritoneal exudate. Any residual erythrocytes were lysed using RBC lysis buffer and the cells were maintained in complete RPMI 1640 media for further experiments.

### Generation of deletion mutant

Δ*chiA* mutant strains were made using one-step deletion strategy as mentioned by Datsenko and Wanner [16]. Briefly, wild-type *Salmonella* (*S.* Typhimurium 14028S or *S*. Typhi CT18) bacteria transformed with a ‘lambda red recombinase’ expressing plasmid under arabinose inducible promoter (pKD46), was grown in LB with 50 µg/ml ampicillin and was induced with 10 mM L-arabinose at 30 °C to an OD_600_ of 0.35-0.4. Electrocompetent cells were prepared by pelleting the bacterial cells and washing the pellet three times with ice cold, sterile MiliQ water and 10% glycerol, followed by resuspension in 50 µl of 10% glycerol. Kanamycin resistance cassette was amplified from pKD4 plasmid using primers containing upstream and downstream sequences of *S.* Typhimurium *chiA* gene (STM14_0022) and *S.* Typhi *chiA* gene (STY0018) fragment. 500 ng of this PCR product was purified and used for electroporation. Transformants were selected on LB agar containing kanamycin plates and were further confirmed with confirmatory primers, *chiA* specific RT primers and kanamycin resistance cassette internal primers.

### Infection and gentamicin protection assay

Epithelial Caco-2 cell line was infected with mid-log phase culture of bacteria grown in LB (OD_600_ 0.3), whereas phagocytic U937 derived monocytes and BMDCs were infected with overnight culture (OD_600_ 0.3). The multiplicity of infection (MOI) of 10 was used in each case. Bacterial attachment to host cells was enhanced by centrifuging at 600 rpm for 10 min. After 25 min of infection, cells were treated with gentamicin (100 μg/ml in complete media) for 1 hour to remove extracellular bacteria and then maintained with 25 μg/ml gentamicin for rest of the experiment. 0.1% Triton-X 100 (v/v in 1x PBS) was used to lyse the cells and the lysate was plated on *Salmonella-Shigella* (SS) agar for *S.* Typhimurium strains and Wilson Blair (WB) agar for *S.* Typhi strains. For invasion assay, cells were lysed after incubation in 100 μg/ml gentamicin treatment (i.e. 1 hour post infection) and percent invasion was calculated with respect to the pre-inoculum used for infection. For intracellular survival assay (ICSA), infected cells were lysed at 2 hours and 18 hours post infection. CFU at 18 hours was divided by CFU at 2 hours to obtain fold replication of the intracellular bacteria. For estimating the cytoplasmic bacterial population, chloroquine resistance assay was performed [32]. Briefly, Caco2 cells were infected by different bacterial strains as mentioned previously. The infected cells were treated with 800 µM chloroquine 1 hour prior to cell lysis and absolute CFU were calculated by plating the cell lysate on selective media.

### Quantitative RT-PCR

Bacterial RNA was isolated from infected cells as described previously by Eriksson *et al.*[11]. Briefly, *Salmonella* infected cells were lysed at different time intervals on ice by incubating for 30 minutes with 0.1% SDS, 1% acidic phenol and 19% ethanol in sterile water. Eukaryotic cell debris was removed by centrifuging the cell lysate at 300g for 10 minutes, followed by pelleting bacterial cells at 5000 rpm for 5 minutes. At each time point, bacteria were recovered from a 6-well plate of infected U937-derived monocytes and pooled to isolate RNA. *In vitro* grown bacterial RNA was obtained by growing bacteria statically at 37 °C in RPMI 1640 medium, under 5 % CO_2_. The bacterial pellet was resuspended in TRIzol reagent (Takara) and stored at −80 °C. Young adult hermaphrodites were infected with respective bacterial strains for 48 hr. Infected worms were harvested by washing the plates with M9 buffer and pelleting at 1000g for 1 min. The extracellular bacteria were removed by repeatedly washing the pellet for 5-6 times. The worms pellet was resuspended in TRIzol reagent (Takara) and stored at −80 °C. RNA was isolated by phase separation method using chloroform. cDNA was synthesized with reverse transcriptase (GCC Biotech). Quantitative PCR was carried out using SYBR Green Q-PCR kit (Takara). Relative expression with respect to control (*act2* gene) was plotted as fold change. Relative expression with respect to control (16s rRNA gene for bacterial genes and *act2* for *C. elegans* genes) was plotted as fold change.

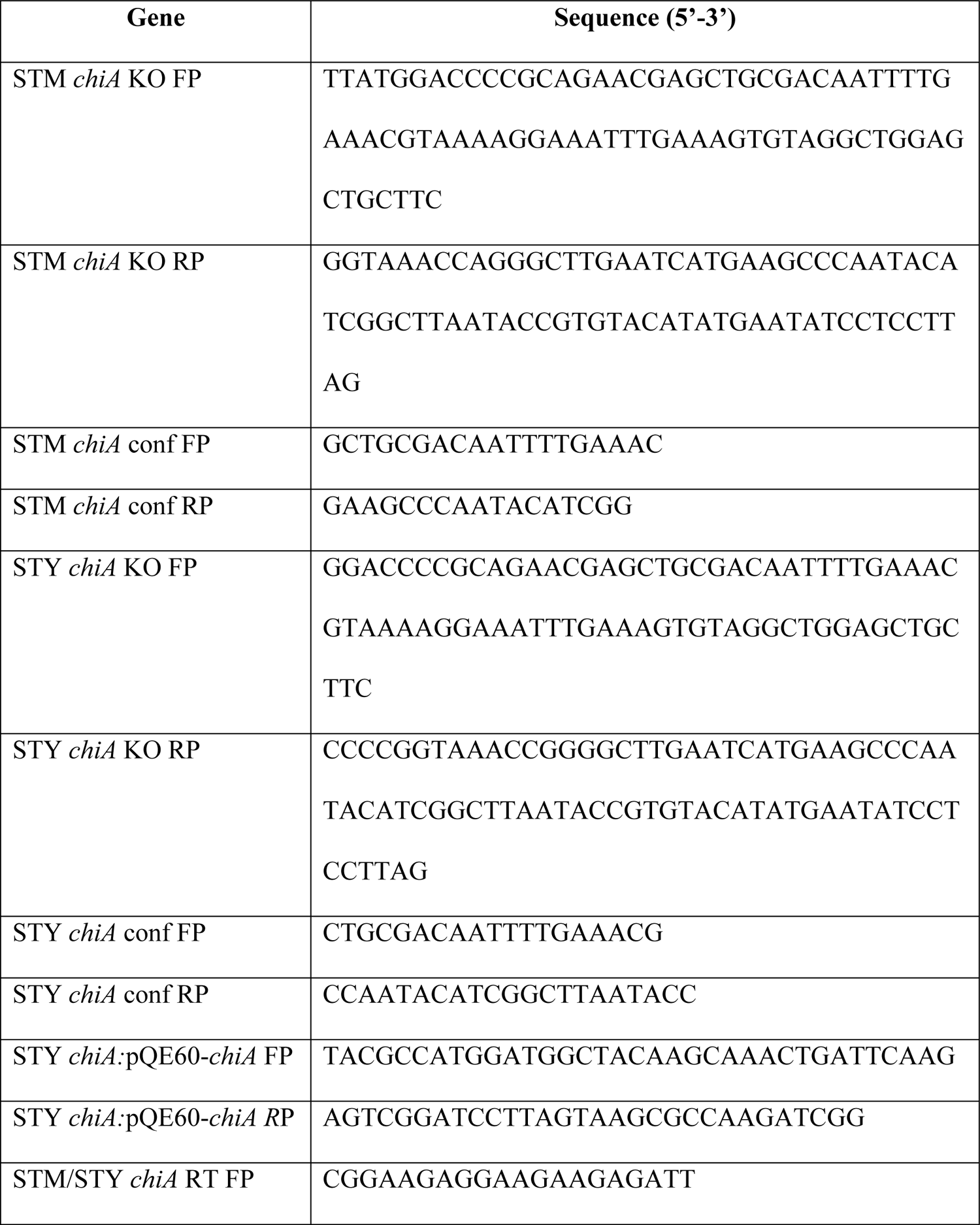
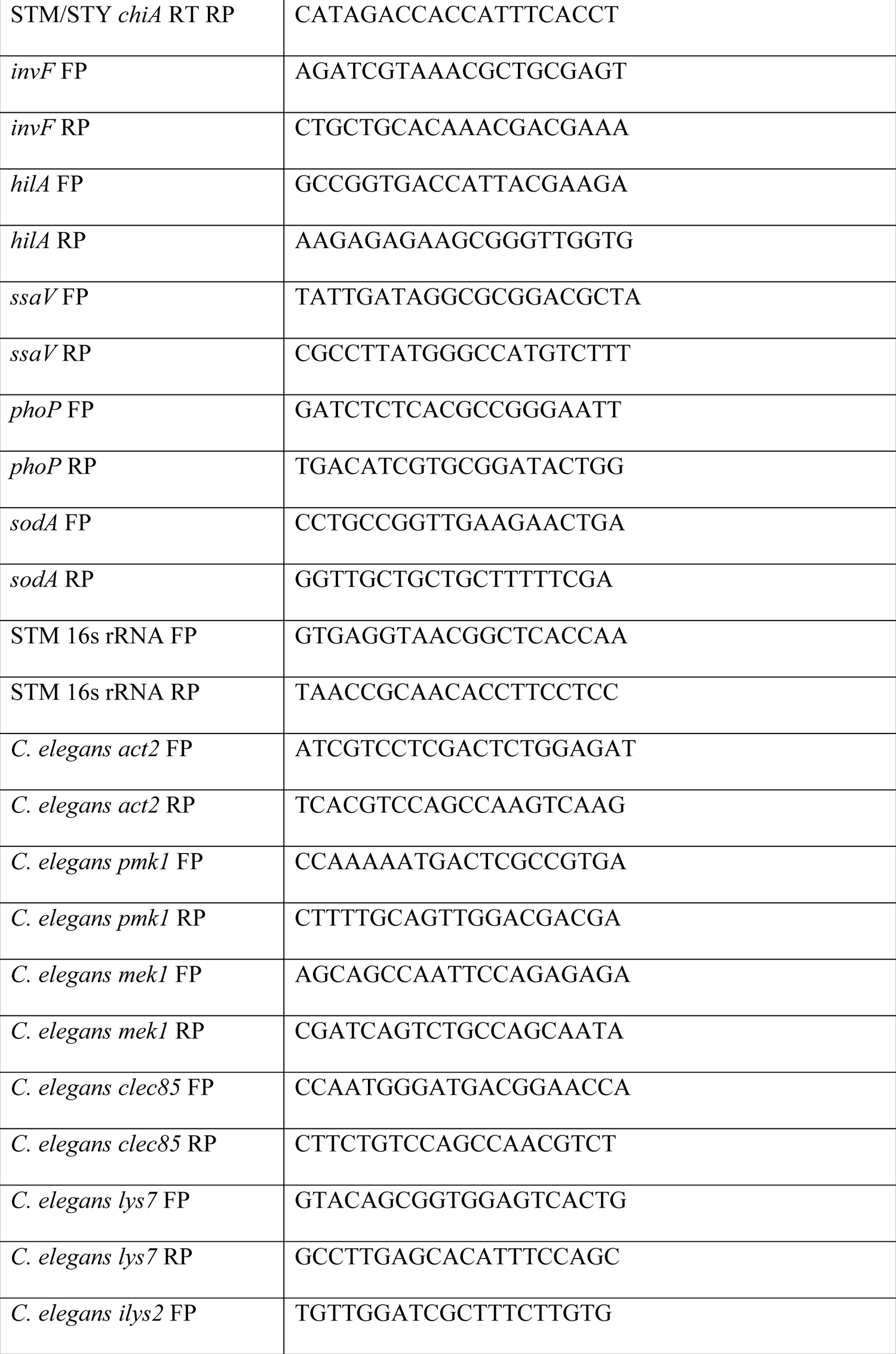
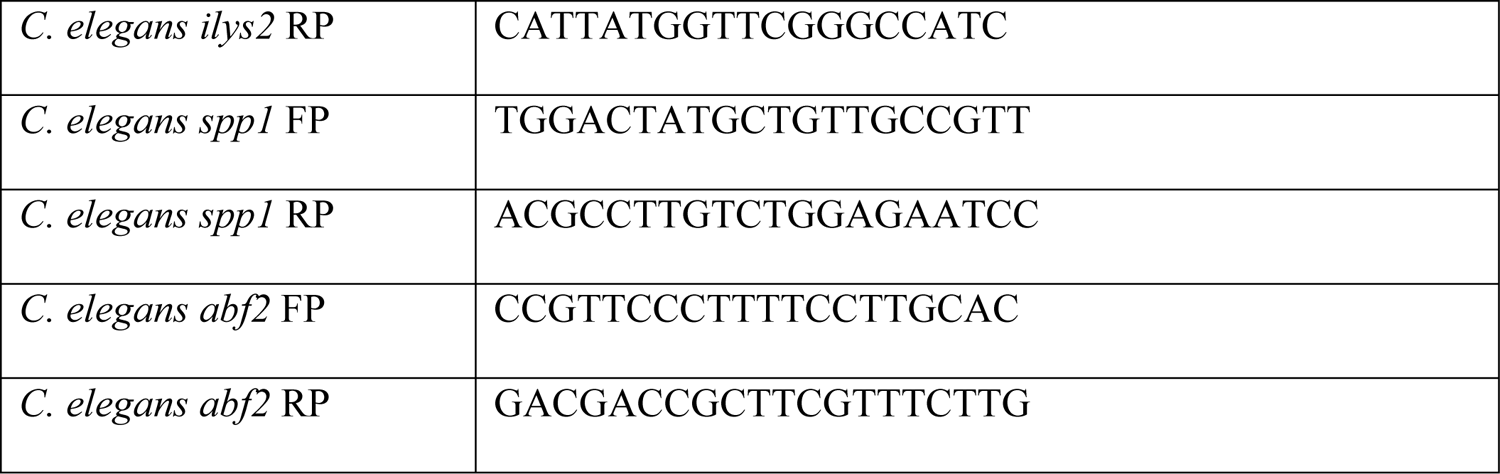
List of strains used in this study.

### Lectin binding assay for cell surface glycan modification

Human colorectal carcinoma cells Caco2 were infected with different bacterial strains as mentioned before. For confocal imaging, cells were seeded on coverslips prior to infection. After infection for the specified time, the cells were fixed with 3.5% PFA for 20 min on ice. For flow cytometry, cells were washed with PBS and treated with 1x Trypsin-EDTA (TE) for 15 min, under 5% CO_2_ at 37 °C. After the cells were dislodged from the wells, the TE was removed and cells were incubated with 1 ml complete media for 20 min under 5% CO_2_ at 37 °C for recovery. To avoid non-specific lectin binding the cells were treated with blocking buffer (PBS+2% FBS) at RT for 15 min. Specific lectins (50µg/ml lectin solution in blocking buffer for every 10^6^ cells) (Vector Laboratories; #FL-1301, #FL-1071, #FL-1001) were added to each samples and incubated for 30 min at RT, followed by washing with blocking buffer. Cells treated with only FITC dye (Merck; #46950) were used as controls.

### Flow cytometry and immunofluorescence

Cells were fixed with 3.5% PFA for 20 min on ice. All staining except for the surface markers (MHC II, CD4 and CD25), were performed in the presence of permeabilizing agent, 0.01% saponin (Sigma) dissolved in 2.5% BSA containing PBS. Flow cytometry analysis was carried out using BD FACSVerse and BD FACSAria and data were analyzed using BD FACSDiva software. Immunofluorescence images were obtained using Zeiss LSM 710 and/or Zeiss LSM 880. The images were analyzed using ZEN Black 2012 platform. For analysis of activated T cell population (CD4^+^ CD25^+^) from infected mice spleen, splenocytes were isolated from mice that survived through 20 days of infection. Total splenocytes were fixed using 3.5% PFA on ice for 20 min, followed by incubation for 1 hour at RT with fluorophore conjugated antibody cocktail in dark. The cells were washed with 1x PBS and analyzed by flow cytometry. Anti-mouse LAMP1 (DSHB; #1D4B) antibody was used for immunofluorescence microscopy. Anti-mouse I-A/I-E (or MHC II) (clone 2G9) FITC (BD Pharmingen; #553623) antibody was used for immunofluorescence microscopy and flow cytometry. Anti-mouse CD4 FITC (Invitrogen; #11-0041-85), Anti mouse CD25 PE (Invitrogen; #12-0251-82) antibodies were used for flow cytometry.

### Nitric oxide estimation

Sodium nitrite (Sigma) standards of 100 μM, 50 μM, 25 μM, 12.5 μM, 6.25 μM and 3.13 μM were prepared by diluting 0.1 M stock in deionized distilled water. Conditioned media from infected cells were collected after indicated time intervals for estimation of nitrite by Greiss assay [33]. 1% sulphanilamide solution was made in 5% phosphoric acid. To 50 μl of the standards and the samples (in triplicates), 50 μl acidic sulphanilamide was added and incubated at RT, in dark for 10 min. After incubation, 50 μl of 0.1% NED (N-1-naphthylethylene diamine dihydrochloride) solution was added to it and incubated for 10 min in dark at RT. OD_520_ was measured within 30 min of appearance of purple/magenta colored product using TECAN Infinite Pro 200 microplate reader.

### ROS measurement

Intracellular ROS was detected by 2’, 7’-dichlorofluorescin diacetate (H_2_DCFDA; Sigma) staining. Cells were stained with 10µM DCFDA at 37 °C in dark. After 30 min, cells were washed with ice cold PBS and harvested followed by flow cytometry analysis at 495/530 nm in BD FACSVerse.

### T cell proliferation assay

WT BMDCs were infected by incubating the bacteria with DCs for 90min, followed by removal of the bacteria and incubating the infected cells with 25µg/ml gentamicin. Total splenocytes were isolated from the spleen of C57BL/6-Tg (TcraTcrb) 1100Mjb/J mice by mechanical disruption. Erythrocytes were lysed by RBC lysis buffer (Sigma) and cells were maintained in complete RPMI-1640. Finally, non-adherent cells were collected and were used for mixed lymphocyte proliferation assay. The proliferation of the lymphocytes in response to antigen stimuli, was detected by incorporation of the ^3^H_1_ as measured by the scintillation counter.

### *In vivo* experiment

6 weeks old male C57BL/6J mice were used for all the *in vivo* mice experiments. All animal experiments were approved by the Institutional Animal Ethics Committee (CAF/Ethics/670/2019) and the National Animal Care Guidelines were strictly followed. 10^8^ CFUs of overnight grown STM WT and STM Δ*chiA* mutant bacteria were used for oral infection for animal survival assay. The control group was orally administered with sterile 1x PBS. Animals were observed for 20 days for survival and body weight was documented. For *in vivo* invasion, the animals were euthanized after 2 hours of gavage, and the bacterial CFUs in Peyer’s patches (PP) were estimated. To check the bacterial shedding, fecal pellets were collected aseptically from infected cohorts after indicated time. Homogenates were plated on SS agar plates and CFUs were counted. For estimating *in vivo* bacterial burden in different organs, a sublethal dose of 10^7^ CFUs of each bacterial strain were used and bacterial CFUs from liver, spleen. MLN and PP were enumerated after indicated time intervals. Spleens were isolated from the animals after 20 days and the length was measured.

### ELISA for serum cytokines and anti-*Salmonella* IgG

Blood collected from infected animals by cardiac puncture under aseptic conditions, was incubated at RT to facilitate coagulation. Serum was then isolated by centrifugation at 5000 rpm for 10 min at RT and stored at −20 °C for further use. Estimation of serum level of different pro-inflammatory cytokines (IL2 and IFNγ) and anti-inflammatory cytokines (IL10 and IL4) was performed according to the manufacturer’s instructions. Anti-*Salmonella* IgG titer was measured by sandwich ELISA as mentioned previously [34]. Briefly, wells were coated with *Salmonella* LPS (200 ng/well; Sigma) at 4 °C overnight. Next day, LPS was removed and the wells were washed with PBST (PBS+0.05% Tween 20), followed by blocking for 1 hour at RT with 5% FBS in PBS to avoid non-specific binding. After blocking, wells were washed with PBST. The serum samples, diluted in blocking buffer, were added to the wells in triplicates and incubated for 2 hours. Subsequently, wells were washed with PBST and anti-mouse IgG (HRP conjugate) was then added to the wells and incubated for 1 hour at RT. Tetramethylbenzidine (TMB; Sigma) was added and the plate was incubated in dark for 20-30 min. The reactions were stopped with 2 N H_2_SO_4_ and the absorbance was measured at 450 nm.

### In vivo colonization in Caenorhabditis elegans

*C. elegans* var. Bristol worms wildtype strain N2, FT63 [xnIs17; dlg-1::GFP + rol-6(su1006)], and CL2166 [dvls19 III; dvls (pAF15)gst-4p::GFP::NLS III] strains were maintained on NGM media at 20 °C. L4 or Young adult N2 hermaphrodite worms were used for *in vivo* experiments. 10^7^ CFU of different bacterial strains were seeded on NGM plates and grown for 16 hours. Young adult N2 worms were fed at 20 °C with the different bacterial strains for 24 hours or 48 hours to check bacterial colonization in the worms [35]. Bacterial CFU was enumerated by plating worms’ lysate from equal number of infected worms on WB agar plates. Fold change was calculated as the ratio of CFU after 48 hours to CFU after 24 hours. For confocal analysis of the worm gut colonization, FT63 worms were used. mCherry expressing bacterial strains were used to visualize the gut colonization.

Further to check worms survivability, 10^7^ CFUs of overnight grown bacterial strains were seeded on 30 mm dishes containing Brain Heart Infusion (BHI) agar media. ∼30-40 young adult worms were added at the center of each plate and survival was monitored [36]. Animals were transferred to fresh bacterial plates every day for first 5 days and then after every 5 days. The worms were scored as live or dead at regular intervals throughout the course of the assay. Worms were considered dead when they failed to respond to touch stimulus. Chitin-rich organs were visualized using Eosin Y stain. After 24 hours of infection, worms were harvested and washed 5 times with M9 buffer, followed by washing the worms pellet with citrate phosphate buffer (0.2 M Na_2_HPO_4_, 0.1 M potassium citrate, pH 6.0). The worms were resuspended in 500 µL citrate-phosphate buffer and 15 µL of 5 mg/ml eosin Y (in 70% ethanol) was added to it. Tubes were incubated at RT, in dark for 10 minutes, followed by centrifugation at 1000g for 1 min for washing. The supernatant was discarded and the pellet was washed with citrate phosphate buffer 5 times to remove excess eosin Y. Effect of bacterial colonization was determined by infecting CL2166 worms for 48 hours with different strains. CL2166 worms possess oxidative stress inducible GFP. Fluorescence of the infected worms was visualized using Zeiss LSM 880 with Multiphoton mode.

### Bacterial persistence assay in *C. elegans*

Young adult N2 worms were infected as mentioned previously. After 24 hours or 48 hours of infection, the worms were harvested and washed 5 times with M9 buffer. After indicated time, ∼30 worms were mounted for confocal imaging. Rest of the worms were transferred to *E. coli* OP50 plate for further 24 hours. These worms were harvested, washed and imaged as mentioned previously.

### Quantification of pharyngeal pumps

The effect of bacterial colonization on the chitin-rich grinder integrity was determined by counting number of pharyngeal pumps per min. Young adult worms were infected as mentioned previously. After indicated infection time, no. of pharyngeal pumps/min was counted for ∼25 worms from each infected plate.

### Fat estimation by Oil red O

Neutral lipids present in the worms was estimated by Oil Red O (ORO; Sigma) staining [29]. Briefly, solution of Oil Red O was prepared in isopropanol (5 mg/ml) and diluted to 60% in water before use. Synchronized L4 animals were allowed to feed on *E. coli* and STY strains for 48 hr. Worms were harvested in M9 buffer, followed by fixing and permeabilizing using MRWB buffer (160 mM KCl, 40 mM NaCl, 14 mM Na_2_-EGTA, 1 mM spermidine-HCl, 0.4 mM spermine, 30 mM Na-PIPES [Na-piperazine N, N’-bis(2-ethanesulfonic acid); pH 7.4], 0.2% β-mercaptoethanol, 0.2% paraformaldehyde) for 1 hour at RT. The animals were stained with 60% ORO at RT. Excess strain was removed by washing twice with 1x PBST (PBS+0.01% Tween 20). Stained animals were mounted on agar pads.

## Statistical analysis

Data were plotted using GraphPad Prism 6 software. Statistical analysis was performed using Student’s t-test or ANOVA as indicated. The results are expressed as mean ± SEM. p values <0.05 was considered to be significant (p values: ****<0.0001, ***<0.001, **<0.01, *<0.05).

## Acknowledgements

We thank the confocal microscopy facility and real-time facility of Dept. of Microbiology and Cell Biology, IISc. We are thankful to SCh lab in MCB, IISc and VS lab in MRGD, IISc for kindly gifting us the *C. elegans* strains. We thank RB lab in MRDG, IISc for their help with the lectin-binding assay. We also thank our laboratory members for their critical input for the manuscript.

## Author Contribution

KC and DC conceived the study and designed experiments. KC performed experiments, analyzed the data, prepared the figures and wrote the manuscript. DC supervised the work. All the authors read and approved the manuscript.

---

**Fig S1.**
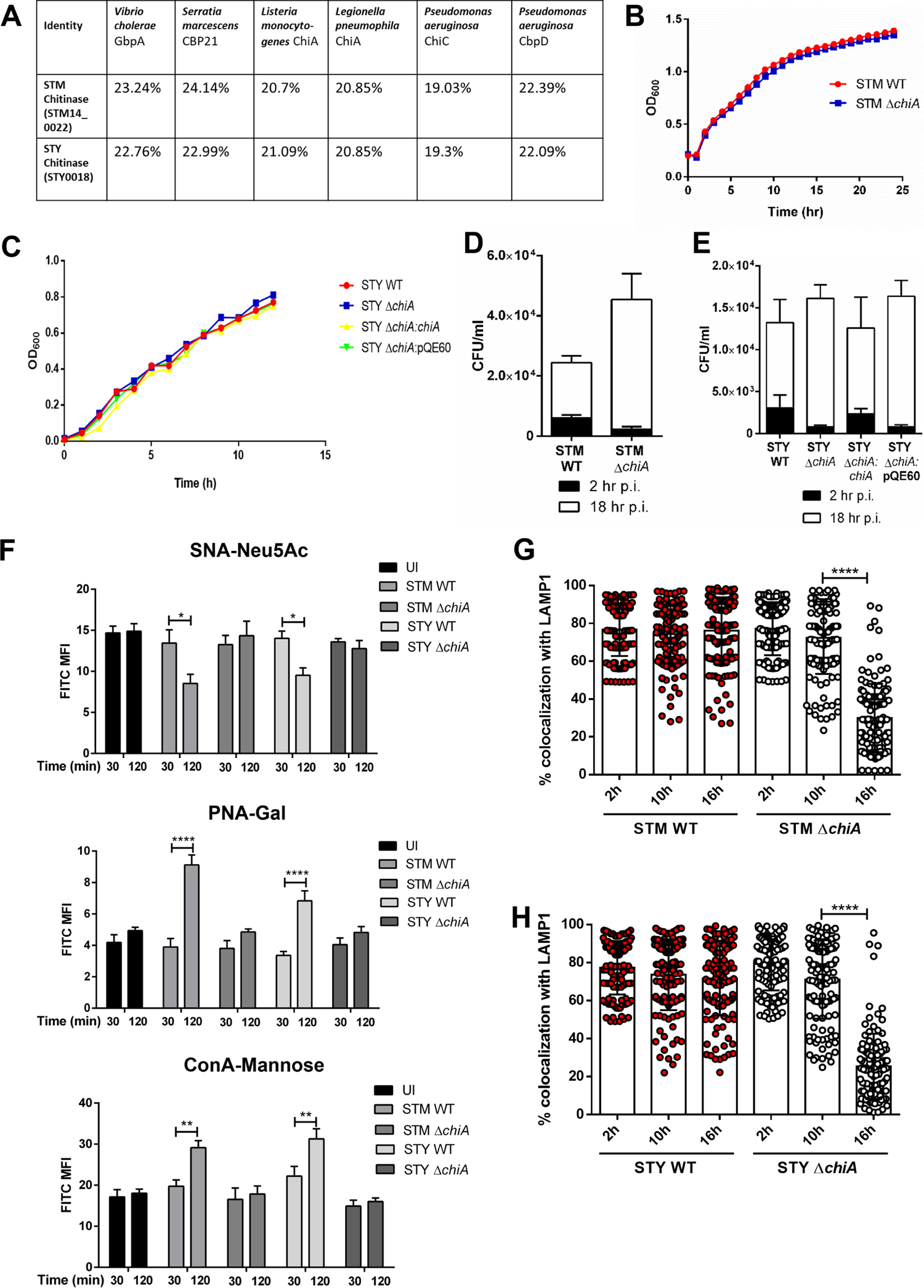

**Fig S2.**
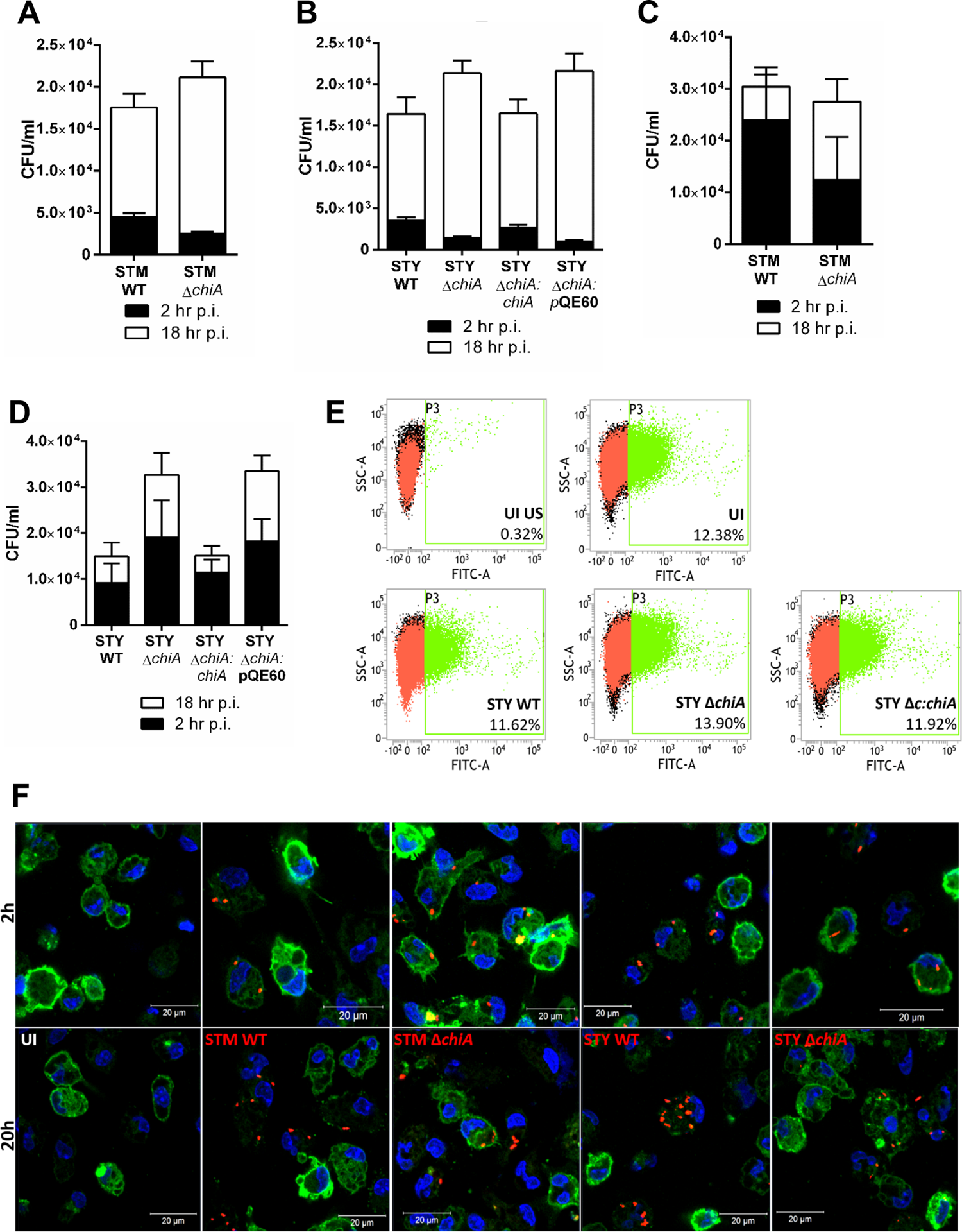

**Fig S3.**
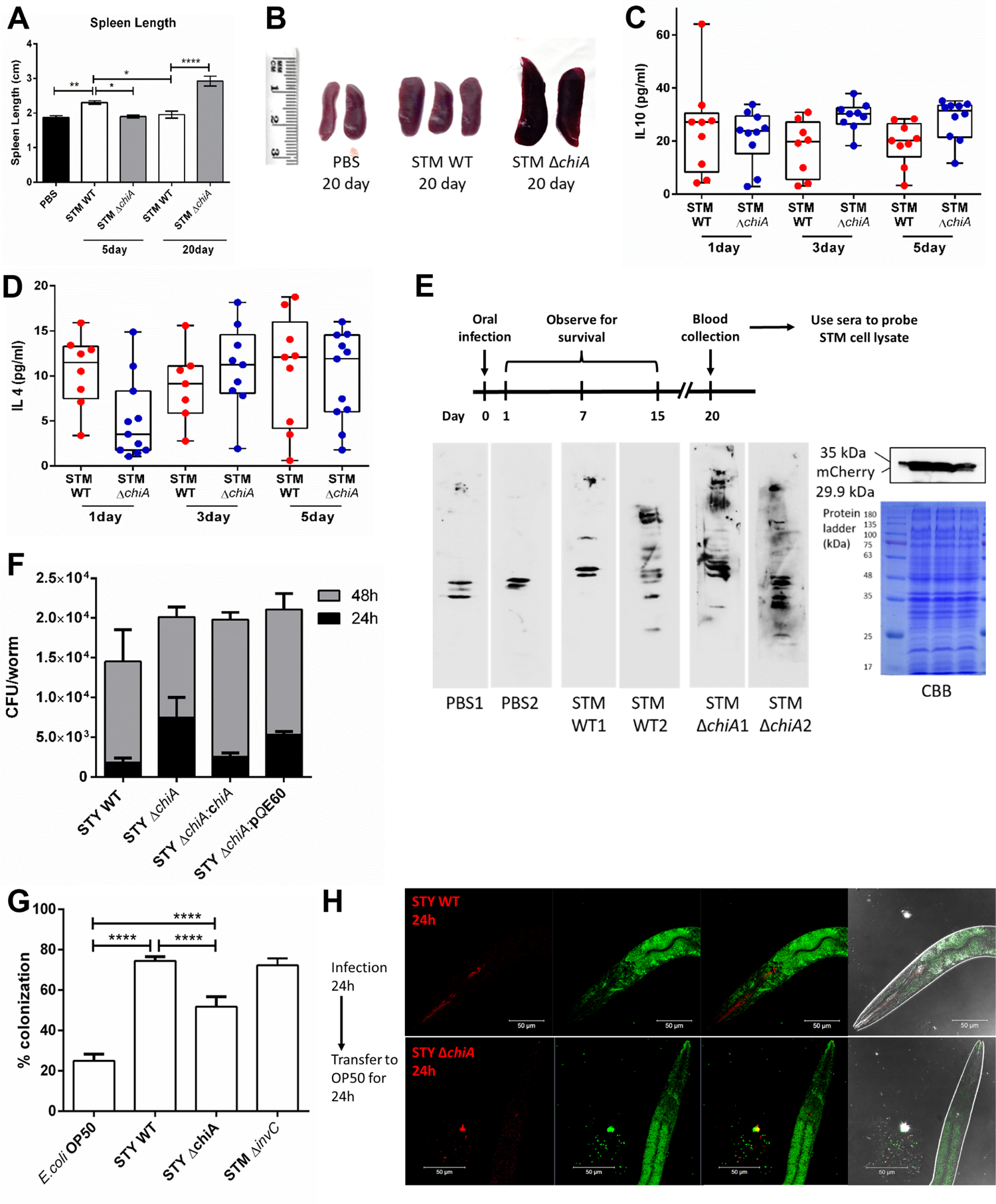

**Fig S4.**
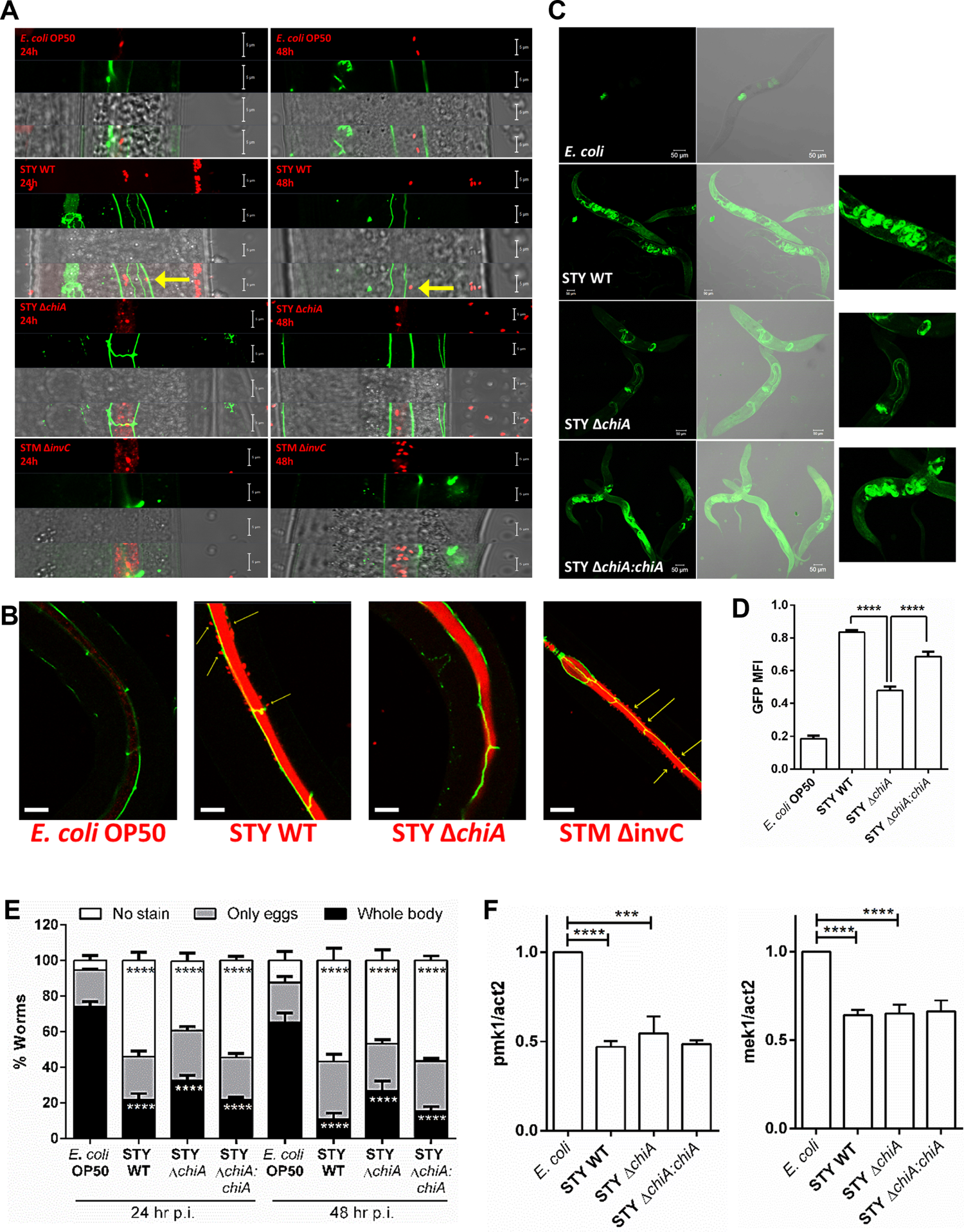

## References

1. Hansson, G.C., Role of mucus layers in gut infection and inflammation. Curr Opin Microbiol, 2012. 15(1): p. 57–62.

2. Frederiksen, R.F., et al., Bacterial chitinases and chitin-binding proteins as virulence factors. Microbiology-Sgm, 2013. 159: p. 833–847.

3. Kirn, T.J., B.A. Jude, and R.K. Taylor, A colonization factor links Vibrio cholerae environmental survival and human infection. Nature, 2005. 438(7069): p. 863-6.

4. Leisner, J.J., et al., Chitin hydrolysis by Listeria spp., including L. monocytogenes. Appl Environ Microbiol, 2008. 74(12): p. 3823–30.

5. Chatterjee, S.S., et al., Intracellular gene expression profile of Listeria monocytogenes. Infect Immun, 2006. 74(2): p. 1323–38.

6. Chaudhuri, S., et al., Contribution of chitinases to Listeria monocytogenes pathogenesis. Appl Environ Microbiol, 2010. 76(21): p. 7302–5.

7. Kawada, M., et al., Chitinase 3-like-1 enhances bacterial adhesion to colonic epithelial cells through the interaction with bacterial chitin-binding protein. Lab Invest, 2008. 88(8): p. 883–95.

8. Salunkhe, P., et al., A cystic fibrosis epidemic strain of Pseudomonas aeruginosa displays enhanced virulence and antimicrobial resistance. J Bacteriol, 2005. 187(14): p. 4908–20.

9. Fung, C., et al., Gene expression of Pseudomonas aeruginosa in a mucin-containing synthetic growth medium mimicking cystic fibrosis lung sputum. J Med Microbiol, 2010. 59(Pt 9): p. 1089–100.

10. DebRoy, S., et al., Legionella pneumophila type II secretome reveals unique exoproteins and a chitinase that promotes bacterial persistence in the lung. Proc Natl Acad Sci U S A, 2006. 103(50): p. 19146–51.

11. Eriksson, S., et al., Unravelling the biology of macrophage infection by gene expression profiling of intracellular Salmonella enterica. Mol Microbiol, 2003. 47(1): p. 103–18.

12. Hautefort, I., et al., During infection of epithelial cells Salmonella enterica serovar Typhimurium undergoes a time-dependent transcriptional adaptation that results in simultaneous expression of three type 3 secretion systems. Cell Microbiol, 2008. 10(4): p. 958–84.

13. Arabyan, N., et al., Salmonella Degrades the Host Glycocalyx Leading to Altered Infection and Glycan Remodeling. Sci Rep, 2016. 6: p. 29525.

14. Park, D., et al., Salmonella Typhimurium Enzymatically Landscapes the Host Intestinal Epithelial Cell (IEC) Surface Glycome to Increase Invasion. Mol Cell Proteomics, 2016. 15(12): p. 3653–3664.

15. Arabyan, N., et al., Implication of Sialidases in Salmonella Infection: Genome Release of Sialidase Knockout Strains from Salmonella enterica Serovar Typhimurium LT2. Genome Announc, 2017. 5(19).

16. Datsenko, K.A. and B.L. Wanner, One-step inactivation of chromosomal genes in Escherichia coli K-12 using PCR products. Proc Natl Acad Sci U S A, 2000. 97(12): p. 6640–5.

17. Bajaj, V., et al., Co-ordinate regulation of Salmonella typhimurium invasion genes by environmental and regulatory factors is mediated by control of hilA expression. Mol Microbiol, 1996. 22(4): p. 703–14.

18. Galan, J.E., Typhoid toxin provides a window into typhoid fever and the biology of Salmonella Typhi. Proc Natl Acad Sci U S A, 2016. 113(23): p. 6338–44.

19. Brumell, J.H., et al., Disruption of the Salmonella-containing vacuole leads to increased replication of Salmonella enterica serovar typhimurium in the cytosol of epithelial cells. Infect Immun, 2002. 70(6): p. 3264–70.

20. Chakravortty, D. and M. Hensel, Inducible nitric oxide synthase and control of intracellular bacterial pathogens. Microbes Infect, 2003. 5(7): p. 621–7.

21. Niedbala, W., B. Cai, and F.Y. Liew, Role of nitric oxide in the regulation of T cell functions. Ann Rheum Dis, 2006. 65 Suppl 3: p. iii37-40.

22. Sato, K., et al., Nitric oxide plays a critical role in suppression of T-cell proliferation by mesenchymal stem cells. Blood, 2007. 109(1): p. 228–34.

23. Cheng, L.E., et al., Enhanced signaling through the IL-2 receptor in CD8+ T cells regulated by antigen recognition results in preferential proliferation and expansion of responding CD8+ T cells rather than promotion of cell death. Proc Natl Acad Sci U S A, 2002. 99(5): p. 3001–6.

24. Vazquez, M.I., J. Catalan-Dibene, and A. Zlotnik, B cells responses and cytokine production are regulated by their immune microenvironment. Cytokine, 2015. 74(2): p. 318–26.

25. Mathur, R., et al., A mouse model of Salmonella typhi infection. Cell, 2012. 151(3): p. 590–602.

26. Labrousse, A., et al., Caenorhabditis elegans is a model host for Salmonella typhimurium. Curr Biol, 2000. 10(23): p. 1543–5.

27. Heustis, R.J., et al., Pharyngeal polysaccharide deacetylases affect development in the nematode C. elegans and deacetylate chitin in vitro. PLoS One, 2012. 7(7): p. e40426.

28. Klein, J.A., et al., Controlled Activity of the Salmonella Invasion-Associated Injectisome Reveals Its Intracellular Role in the Cytosolic Population. mBio, 2017. 8(6).

29. Escorcia, W., et al., Quantification of Lipid Abundance and Evaluation of Lipid Distribution in Caenorhabditis elegans by Nile Red and Oil Red O Staining. J Vis Exp, 2018(133).

30. Steeb, B., et al., Parallel exploitation of diverse host nutrients enhances Salmonella virulence. PLoS Pathog, 2013. 9(4): p. e1003301.

31. Cheminay, C., A. Mohlenbrink, and M. Hensel, Intracellular Salmonella inhibit antigen presentation by dendritic cells. J Immunol, 2005. 174(5): p. 2892–9.

32. Knodler, L.A., V. Nair, and O. Steele-Mortimer, Quantitative assessment of cytosolic Salmonella in epithelial cells. PLoS One, 2014. 9(1): p. e84681.

33. Green, L.C., et al., Analysis of nitrate, nitrite, and [15N]nitrate in biological fluids. Anal Biochem, 1982. 126(1): p. 131–8.

34. Datey, A., et al., Rewiring of one carbon metabolism in Salmonella serves as an excellent live vaccine against systemic salmonellosis. Vaccine, 2018. 36(50): p. 7715–7727.

35. Tan, M.W., S. Mahajan-Miklos, and F.M. Ausubel, Killing of Caenorhabditis elegans by Pseudomonas aeruginosa used to model mammalian bacterial pathogenesis. Proc Natl Acad Sci U S A, 1999. 96(2): p. 715–20.

36. Everman, J.L., et al., Establishing Caenorhabditis elegans as a model for Mycobacterium avium subspecies hominissuis infection and intestinal colonization. Biol Open, 2015. 4(10): p. 1330–5.

